# High-resolution analysis of the treated coeliac disease microbiome reveals strain-level variation

**DOI:** 10.1101/2024.03.08.584098

**Authors:** Jelle Slager, Hanna L. Simpson, Ranko Gacesa, Lianmin Chen, Ineke L. Tan, Jody Gelderloos, Astrid Maatman, Cisca Wijmenga, Alexandra Zhernakova, Jingyuan Fu, Rinse K. Weersma, Gieneke Gonera, Iris H. Jonkers, Sebo Withoff

## Abstract

**Background:** Coeliac disease (CeD) is an immune-mediated disorder primarily affecting the small intestine, characterised by an inflammatory immune reaction to dietary gluten. CeD onset results from a multifaceted interplay of genetic and environmental factors. While recent data show that alterations in gut microbiome composition could play an important role, many current studies are constrained by small sample sizes and limited resolution.

**Methods:** To address these limitations, we analysed faecal gut microbiota from two Dutch cohorts, CeDNN (128 treated CeD patients (tCeD), 106 controls) and the Lifelines Dutch Microbiome Project (24 self-reported tCeD, 654 controls), using shotgun metagenomic sequencing. Self-reported IBS (570 cases, 1710 controls) and IBD (93 cases, 465 controls) were used as comparative conditions of the gastrointestinal tract. Interindividual variation within the case and control groups was calculated at whole microbiome and strain level. Finally, species-specific gene repertoires were analysed in tCeD patients and controls.

**Results:** Within-individual microbiome diversity was decreased in patients with self-reported IBS and IBD but not in tCeD patients. Each condition displayed a unique microbial pattern and, in addition to confirming previously reported microbiome associations, we identify an increase in the levels of *Clostridium sp. CAG:253*, *Roseburia hominis*, and *Eggerthella lenta*, amongst others. We further show that the observed changes can partially be explained by gluten-free diet adherence. We also observe increased interindividual variation of gut microbiome composition among tCeD patients and a higher bacterial mutation frequency in tCeD that contributes to higher interindividual variation at strain level. In addition, the immotile European subspecies of *Eubacterium rectale*, which has a distinct carbohydrate metabolism potential, was nearly absent in tCeD patients.

**Conclusion:** Our study sheds light on the complex interplay between the gut microbiome and CeD, revealing increased interindividual variation and strain-level variation in tCeD patients. These findings expand our understanding of the microbiome’s role in intestinal health and disease.

**Highlights:**

- This is the largest microbiome collection of coeliac disease (CeD) patients assembled to date, providing insights into gut microbiome composition down to strain level.
- Compared to controls, treated CeD (tCeD) patients adhering to a gluten-free diet show novel gut microbiome associations.
- tCeD patients also have a less uniform gut microbiome than controls.
- Bacteria display higher mutation frequency in tCeD compared to controls.
- Hallmarks of the European subspecies of *Eubacterium rectale* are nearly absent in tCeD patients, implying selection at strain level.

## Introduction

Coeliac disease (CeD) is a multifactorial autoimmune disease characterised by an inflammatory response in the small intestine^1^. It arises in genetically susceptible individuals carrying HLA-DQ2 or -DQ8 haplotypes and is triggered by the ingestion of dietary gluten, a storage protein found in grains such as wheat, barley, and rye^1–3^. CeD is relatively common, affecting ∼1–2% of the global population^4,5^. In CeD pathogenesis, an inflammatory immune response is initiated by the translocation of partially digested gluten peptides from the intestinal lumen into the lamina propria. The gluten peptides undergo deamidation by tissue transglutaminase 2 (tTG2), which confers increased binding affinity to class II HLA-DQ2 or - DQ8 molecules on antigen-presenting cells^6^. Notably, tTG2 activity has also been reported in the intestinal lumen, suggesting that deamidation may also occur before transepithelial passage of the peptides^7^. Presentation of deamidated gluten peptides by antigen-presenting cells leads to activation of gluten-specific CD4^+^ T cells^8–10^. A complex cytokine-mediated cascade then ensues, ultimately resulting in the destruction of the epithelial barrier by intraepithelial lymphocytes^11^. Currently, the only treatment option for CeD is strict lifelong adherence to a gluten-free diet (GFD), which necessitates extensive patient education and ongoing monitoring^12^.

CeD presents a unique case in which the trigger and major contributing genetic factors are known yet not all HLA-DQ2- or -DQ8-positive individuals develop CeD, suggesting that other non-genetic factors contribute to its development^13–16^. One such factor is the gut microbiome, and multiple studies have reported changes in the composition and function of the gut microbiome in CeD. These include reduced levels of beneficial bacteria such as *Bifidobacterium* and *Lactobacillus* species and increased levels of *Proteobacterium* (renamed to *Pseudomonadota*^17^) species, compared to healthy controls^18–22^. Changes in gut microbiome composition have also been reported to precede CeD development^23,24^.

Several mechanisms have been proposed that implicate changes in microbiome composition in CeD pathophysiology. Specific microbial proteases have been reported to be involved in the differential processing of gluten peptides, altering their immunogenicity and thus their ability to incite an inflammatory immune response^25–27^. Moreover, toxins and other metabolites secreted by pathobionts may directly or indirectly affect intestinal barrier permeability, influencing the passage of gluten peptides into the lamina propria^28–30^. In addition, microbial proteins containing sequences similar to immunogenic gluten peptides (molecular mimicry) may cause a similar immune response^31^. Lastly, given the gut microbiome’s important role in the development and homeostasis of the mucosal immune system, alterations in its composition may contribute to an overall loss of tolerance to gluten. While the microbiome is often implicated in CeD onset, it may also undergo changes as a consequence of the disease, driven by factors such as the inflammatory milieu in the small intestine or the increased epithelial permeability observed in CeD patients, ultimately leading to persistent dysbiosis^32,33^. Altogether, these findings underscore that there is an intricate interplay between gut microbiota, gluten, and the immune system in CeD development. Although previous studies investigating the gut microbiome of CeD patients have provided important insights, many have been constrained by small patient cohorts^23,24,26,34,35^, which limits statistical power, and/or relied on 16S sequencing^26,34–36^, which limits the taxonomic resolution to genus level. Furthermore, most studies do not provide information on gut microbiota functionality.

To overcome these issues, in this study we used metagenomic sequencing (MGS) to profile the microbiome in the largest CeD cohort to date, providing subspecies-level resolution. In addition to confirming previous findings^37,38^, our results reveal novel associations between treated CeD (tCeD) and the microbiome, the contribution of a GFD to gut microbiome composition, and a less uniform microbiome composition with increased strain-to-strain variation in patients with tCeD. In our taxonomic analyses, we demonstrate that tCeD and self-reported inflammatory bowel disease (IBD) show unique microbiome signatures with limited overlap. Lastly, we report shifts in the subspecies distribution of *Eubacterium rectale* and *Firmicutes bacterium CAG:83* among tCeD patients compared to controls.

## Materials and Methods

### Study population

#### Treated CeD

The Celiac Disease Northern Netherlands cohort (CeDNN, currently unpublished) comprises 128 CeD patients and 106 controls who provided faecal samples at project onset, with complete metadata including age and sex. Most CeDNN patients had already received a clinical diagnosis and were adhering to a GFD at the time of inclusion. Since nearly all controls were relatives or partners of the included patients, i.e. sharing genetic and lifestyle factors, we enhanced our study by merging this dataset with a subset of participants from the Lifelines Dutch Microbiome Project (LL-DMP)^40^. From the LL-DMP, we included 24 self-reported CeD patients and 654 controls. Controls were selected such that the combined dataset contained five matched (age, sex, sequencing depth) controls for each CeD patient (Table 1). Propensity score matching was performed with the R program *MatchIt* (v4.5.5)^41^, and self-reported IBD and irritable bowel syndrome (IBS) patients were excluded from the control group. As only 9 of the 152 (6%) CeD patients included reported not adhering to a GFD, our study group is best described as a tCeD cohort.

**Table 1:** Description of the study population.

| Variable | Cohort | CeD |  | IBD |  | IBS |  |
| --- | --- | --- | --- | --- | --- | --- | --- |
|  |  | CeD | Controls | IBD | Controls | IBS | Controls |
| Sample size | CeDNN | 128 | 106 |  |  |  |  |
|  | LL-DMP | 24 | 654 | 93 | 465 | 570 | 1710 |
| Mean age (range) |  | 30 (3–82) | 30 (2–82) | 53 (33–74) | 54 (13–81) | 49 (19–84) | 49 (13–83) |
| Sex female (%) |  | 97 (64%) | 445 (59%) | 56 (60%) | 267 (57%) | 411 (72%) | 1233 (72%) |
| Mean BMI (SD) |  | 19.0 (5.1)<sup>a</sup> | 21.7 (4.8) | 25.9 (3.9) | 25.9 (3.9) | 25.8 (4.4) | 25.8 (4.3) |
| Gluten-free diet |  | 94% | 5%<sup>b</sup> | 0%<sup>b</sup> | 0%<sup>b</sup> | 0.2%<sup>b</sup> | 0.2%<sup>b</sup> |

Summary statistics of the data used in this study. <sup>a</sup>BMI of adult patients (UMCG) was not available from medical records, skewing the BMI data of CeD patients (median age 12) relative to controls (median age 16), which explains most of the observed difference. <sup>b</sup>Estimation based on a subpopulation of Lifelines participants who filled out the relevant food frequency questionnaire<sup>39</sup>. CeD, coeliac disease. IBD, inflammatory bowel disease. IBS, irritable bowel syndrome.

#### IBD and IBS

For comparative conditions, 93 self-reported IBD patients (Crohn’s disease or ulcerative colitis) from the LL-DMP were each matched with five controls (based on age, sex, sequencing depth; 465 controls total) and 570 self-reported IBS patients from LL-DMP were each matched with three controls (age, sex, sequencing depth; 1710 controls total). Self-reported CeD, IBD, and IBS patients were excluded from both control groups (Table 1).

### Ethical approval

The LL-DMP study and the CeDNN study were approved by the medical ethical committee of the University Medical Center Groningen (METc numbers 2017/152 and 2013/440, respectively). Additional written consent was signed by all participants, their parents, or legal representatives (for participants aged under 16 for CeDNN and under 18 for LL-DMP).

### Phenotypic data

CeDNN uses the same questionnaires as Lifelines. Clinical information was available for many participants, including date of diagnosis and adherence to a GFD. The time between diagnosis and faecal sampling served as a proxy for the duration of GFD adherence.

### Faecal sample collection, DNA extraction, and sequencing

Faecal sample collection, DNA extraction, and metagenomic sequencing were performed as previously described^40^. In summary, participants from both CeDNN and LL-DMP provided stool samples from home. Participants were asked to freeze stool samples within 15 minutes of production. Frozen samples were collected by personnel/students of the University Medical Center Groningen (CeDNN) or Lifelines (LL-DMP), transported to their respective biorepositories on dry ice, and stored at −80 °C until DNA extraction. Microbial DNA was isolated with the QIAamp Fast DNA Stool Mini Kit (QIAGEN), according to the manufacturer’s instructions, using the QIAcube (QIAGEN) automated sample preparation system. Library preparation for samples with total DNA yield <200 ng (determined by Qubit 4 Fluorometer) was performed using the NEBNext Ultra DNA Library Prep Kit for Illumina. Libraries for other samples were prepared using the NEBNext Ultra II DNA Library Prep Kit for Illumina. Metagenomic sequencing was performed at Novogene, China using the Illumina HiSeq 2000 platform to generate approximately 8 Gb of 150 nucleotide paired-end reads per sample.

### Metagenomic data processing

#### Trimming and filtering of reads

Metagenomes were profiled as previously stated in the analyses of the 1000IBD^42^, Lifelines-DEEP^43^, and Lifelines-DMP cohorts^40^. lllumina adapters and low-quality reads (Phred score <30) were filtered out using *KneadData* (v0.10.0 with trimmomatic options: LEADING:20 TRAILING:20 SLIDINGWINDOW:4:20 MINLEN:50)^44^. Following trimming, the KneadData-integrated *Bowtie2* tool (v2.5.0)^45^ was used to remove reads aligned to the human genome (GRCh37/hg19).

#### Taxonomy and pathway analysis

The taxonomic composition of metagenomes was profiled with *MetaPhlAn3* (v3.0.1)^46^ using the MetaPhlAn database of marker genes *mpa_v30*. None of the included samples had a eukaryotic or viral abundance >25% of total microbiome content or a total read depth <10 million. MetaCyc pathway profiling was performed with *HUMAnN3* (v3.0.1) with default settings^46^ integrated with the *DIAMOND* alignment tool (v2.0.15)^47^, UniRef90 protein database (v0.1.1)^48^, and *ChocoPhlAn* pan-genome database (*ChocoPhlAn_201901*)^46^. Since individual genes can be mapped to multiple pathways, the pathway abundances were instead normalised to the sum of gene family abundances per sample.

To preserve statistical power, bacterial taxa and pathways were excluded from further analyses if present at a minimum abundance of 0.001% in <5% (taxa) or <2.5% (pathways) of samples. This yielded 301 taxa (6 phyla, 14 classes, 20 orders, 33 families, 70 genera, and 158 species) and 338 pathways for tCeD vs. controls, 310 taxa (7 phyla, 15 classes, 20 orders, 31 families, 71 genera, and 166 species) and 349 pathways for IBD vs. controls, and 316 taxa (9 phyla, 17 classes, 23 orders, 35 families, 73 genera, and 159 species) and 331 pathways for IBS vs. controls.

#### UpSet plot

To visualise the overlap among observed differences in the three comparisons (tCeD, IBD, or IBS vs. their respective matched controls), data was plotted using the *UpSetR* package in R^49^.

#### Alpha and beta diversity and interindividual variation

Based on the abundance profiles of the taxa that passed the filtering process, we calculated alpha diversity, as measured by Shannon entropy, at genus- and species-level using the *diversity* function in R package *vegan* (v.2.6.4)^50^. Beta diversity was assessed through Bray-Curtis dissimilarity indices determined with the *vegdist* function from R package *vegan*.

#### Strain- and gene-level analyses

For the comparison between individuals with tCeD and controls, we focused on differences at a subspecies level. Initially, we conducted strain-level profiling using StrainPhlAn3 (v3.0)^46^, starting with default parameters for every species. For the species where we found the default parameters to be overly stringent, but that were significantly associated with CeD according to the MetaPhlAn output, we then adjusted the StrainPhlAn3 parameters to, subsequently: “--marker_in_n_samples 50” and “--marker_in_n_samples 20 --sample_with_n_markers 10”.

Among these adjustments, three species (*Eisenbergiella tayi*, *Haemophilus parainfluenzae*, and *Anaerotruncus colihominis*) were only successfully analysed with the most lenient parameter set. Only *Enterococcus faecium* failed with all attempted settings. Distance matrices were constructed from the multiple alignments using the command line tool *distmat*. We then used *PanPhlAn3*^46^ to analyse the gene repertoire of the seven species that were detected in at least 10 tCeD cases and 10 controls and showed at least a nominally significant association between analysis group and interindividual strain distance or strain cluster distribution (see partition around medoids (PAM) clustering below). Heatmaps were generated with the *ComplexHeatmap* package in R^51^. Analyses were performed using locally installed tools and databases on the high-performance computing infrastructure available at our institution using the MOLGENIS data platform^52^.

### Statistical analyses

#### Associations between alpha diversity and sample group

We conducted a comparison of Shannon diversity between disease groups (tCeD, IBD, and IBS) and controls using a linear model. Specifically, when comparing tCeD to controls, we included sequencing batch (CeDNN_batch1, CeDNN_batch2, and LL-DMP) as a covariate. No significant batch effects were observed for alpha diversity.

#### Associations of sample group with bacterial taxa and MetaCyc pathways

We tested for differentially abundant bacterial taxa and MetaCyc pathways by modelling their abundance using linear models with FDR-correction. Differences in prevalence between sample groups were tested using logistic regression, followed by a χ^2^-test using the *step* function from the R package *stats* (with FDR-correction). For the comparisons of tCeD versus controls, sequencing batch (CeDNN_batch1, CeDNN_batch2, and LL-DMP) was used as a covariate. The correlation between taxonomic and pathway abundance was quantified using Kendall’s τ coefficient. As sequencing batch was to some extent confounded with disease status (most included patients are from CeDNN, whereas most controls are from LL-DMP), we indicate which associations with disease status also showed a significant batch effect.

#### Associations between GFD duration and bacterial abundance

The association of bacterial abundance with GFD duration was assessed within the group of CeD patients from the CeDNN cohort, exclusively for bacterial species with significantly higher abundance in tCeD than in controls. GFD duration was set to 0 months for participants not on a GFD. Using a linear model, the effects of age, sex, sequencing depth, and sequencing batch on bacterial abundance were regressed. Spearman’s ρ tests were then used to assess the effect of GFD duration on the residuals of bacterial abundance. We determined confidence intervals with the *SpearmanRho* function from the *DescTools* package^53^ and p-values with the *cor.test* function from the *stats* package.

Separately, 11 controls from LL-DMP reporting GFD adherence were each matched with five controls (based on age, sex, sequencing depth; 55 controls total) that reported not to adhere to any diet. Self-reported CeD and IBD patients were excluded from both groups. Testing for differential abundance and prevalence in participants adhering to a GFD was performed as described for comparisons between patients and control.

#### Beta diversity

To evaluate the relationship between microbiome composition and disease groups, we conducted a PERMANOVA test using the *adonis2* function from the *vegan* package^50^. We utilised Bray-Curtis distance metrics and performed 999 permutations for each test. When comparing tCeD to controls, sequencing batch (CeDNN_batch1, CeDNN_batch2, and LL-DMP) was incorporated as a covariate. No significant batch effects were observed on beta diversity.

To measure the heterogeneity within each sample group (tCeD, IBD, IBS, and their respective control groups), we computed the distribution of all pairwise Bray-Curtis distances, excluding sample pairs related to each other. Significance was determined using permutation tests with 1000 permutations.

#### Strain-level analyses

To investigate whether increased interindividual differences occur at strain level, we measured strain-to-strain variation using the distance matrices generated by *distmat*. These matrices served as proxies for the mutational distance between strains of the same species in different samples within either the control or tCeD group. Sample pairs that were related were excluded from this analysis. Significance was assessed using permutation tests with 1000 permutations.

We then conducted PAM clustering with the *pam* function from the *cluster* package in R^54^. The *prediction.strength* function from the *fpc* package (*clustermethod=claraCBI*) was used to determine the number of clusters that best described the data^55^. Correlation between tCeD and strain clusters was examined using Fisher’s exact tests, with FDR-correction for multiple comparisons.

#### Gene prevalence and gene enrichment analysis

Associations between the prevalence of UniRef90 gene families, as determined by *PanPhlAn*, and tCeD were determined using logistic regression, followed by a χ^2^-test using the *step* function from the *stats* package in R (with FDR-correction). Age, sex, sequencing depth, and sequencing batch were used as covariates. Genes present in <20% or >80% of all samples were excluded from this analysis. Protein family (Pfam) and Gene Ontology annotations were added to UniRef90 gene families, and gene enrichment analysis was performed using Fisher’s exact tests with FDR-correction. Nominally significant gene families were used as input for the enrichment tests. No significant batch effects were observed for any genes with a significant association with tCeD.

#### Correlation between RadC-like JAB domain proteins and interindividual strain distance

To investigate whether the presence of proteins with the RadC-like JAB domain (PF04002), as identified by PanPhlAn, influences the mutational load of associated bacterial species (*B. ovatus*, *B. uniformis*, *B. xylanisolvens,* and *B. vulgatus*), we conducted the following analysis. For each sample analysed by both StrainPhlAn and PanPhlAn, we recorded the median mutational distance (determined by StrainPhlAn) to all other samples within the same sample group (either tCeD or control) and the number of proteins with a RadC-like JAB domain identified by PanPhlAn. We then assessed trends within sample groups using a linear model. We also examined any negative trend, irrespective of the sample group, using a linear mixed-effects model (implemented with the *lmer* function from the *lmerTest* package^56^ in R), with the sample group as a random effect.

## Results

### Alpha diversity is decreased in self-reported IBS and IBD but not in tCeD

Although contradictory data exist in literature, it is most commonly reported that the within-sample microbiome diversity (alpha diversity) in untreated CeD is higher than that in controls^37,57^ and that adherence to a GFD restores alpha diversity to pre-disease levels^37,38^. Indeed, using the Shannon index to assess microbiome richness and evenness, we did not observe a significant difference in alpha diversity on genus-(1.84 in tCeD vs. 1.86 in controls) or species-level (2.48 tCeD vs. 2.44 controls). Self-reported IBD and IBS patients did display a small but significant (p < 0.05) decrease in alpha diversity at genus level compared to matched controls (1.70 IBD vs. 1.87 controls, 1.83 IBS vs. 1.87 controls), which is in line with studies that observed a marked decrease in diversity, particularly in more inflamed IBD-like samples^58–60^. Although tCeD patients showed comparable alpha diversity to controls, we identified several differences when studying the microbiome composition in more detail.

### tCeD, IBD, and IBS show distinct microbial patterns

Next, we analysed the abundance and prevalence of faecal bacteria in individuals with tCeD (Supp. Table S1), IBD (Supp. Table S2), and IBS (Supp. Table S3), along with their matched control groups. Our findings revealed strikingly different trends among these sample sets (Table 2, Figure 1). While all three disease groups were characterised by an increase in several *Bacteroides* species compared to controls, we saw notable differences within the genus (p_adj_ < 0.05 in one condition, nominal p > 0.05 in other conditions). For example, *B. intestinalis*, *B. stercoris,* and *B. uniformis* were exclusively altered in IBD, whereas *B. xylanisolvens* was only enriched in tCeD. Only *B. ovatus* was enriched in more than one subject group (tCeD and IBD). All other species with changed abundance in tCeD were unaltered in IBD and IBS. Only three taxa (*Bacteroides* and *Veillonella* genera and *P. merdae*) showed significant differential abundance in self-reported IBS patients compared to controls, in line with earlier studies (see review by Pittayanon et al.^61^). Although differences in prevalence were less evident between diseases (Table 3), the prevalence of several bacteria was exclusively affected in tCeD. Together, these results suggest that changes in the microbiome are disease-specific rather than being primarily driven by the overall inflammatory intestinal milieu or barrier defect.

**Figure 1:**
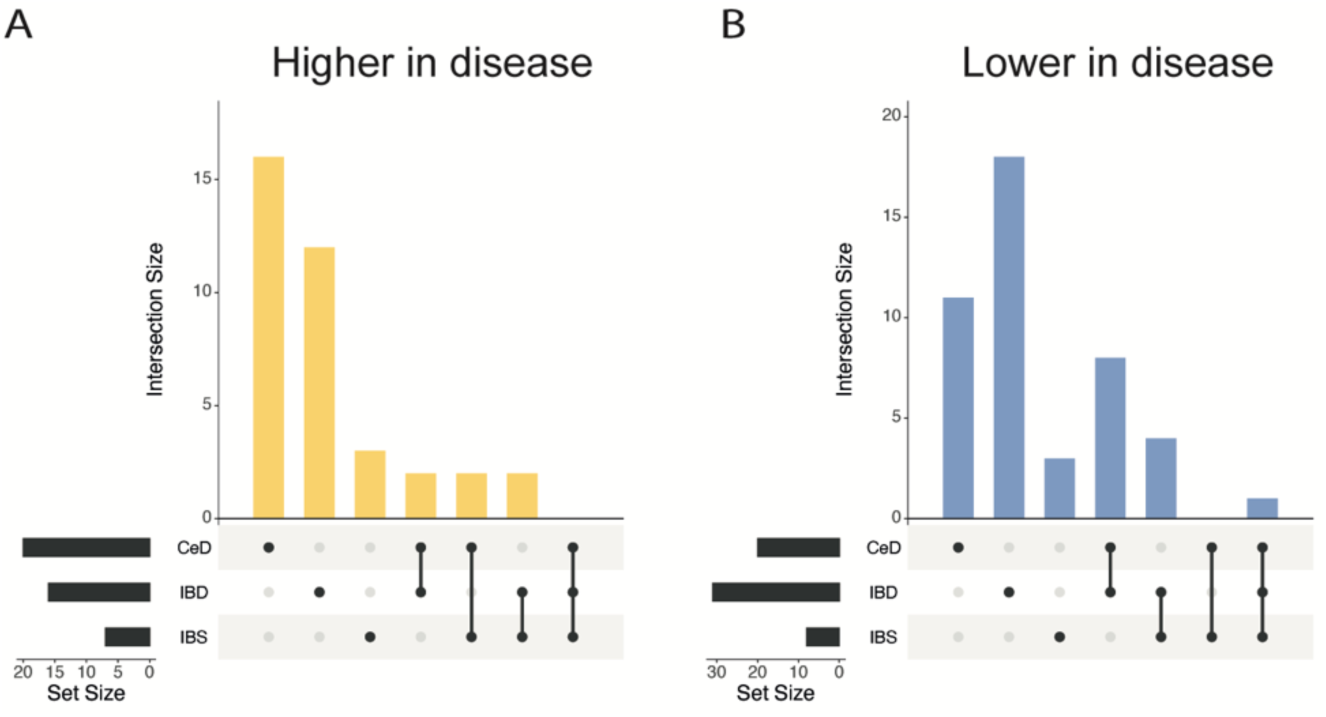
Overlap in microbiome changes between treated coeliac disease (tCeD), inflammatory bowel disease (IBD), and irritable bowel syndrome (IBS). Bacterial species with (A) higher abundance or prevalence in disease and (B) lower abundance or prevalence in disease. Vertical bars indicate the number of species affected in the disease groups, as indicated by the black dots below each respective bar. Horizontal black bars at bottom left indicate the total number of species affected in each disease group.

**Table 2:**
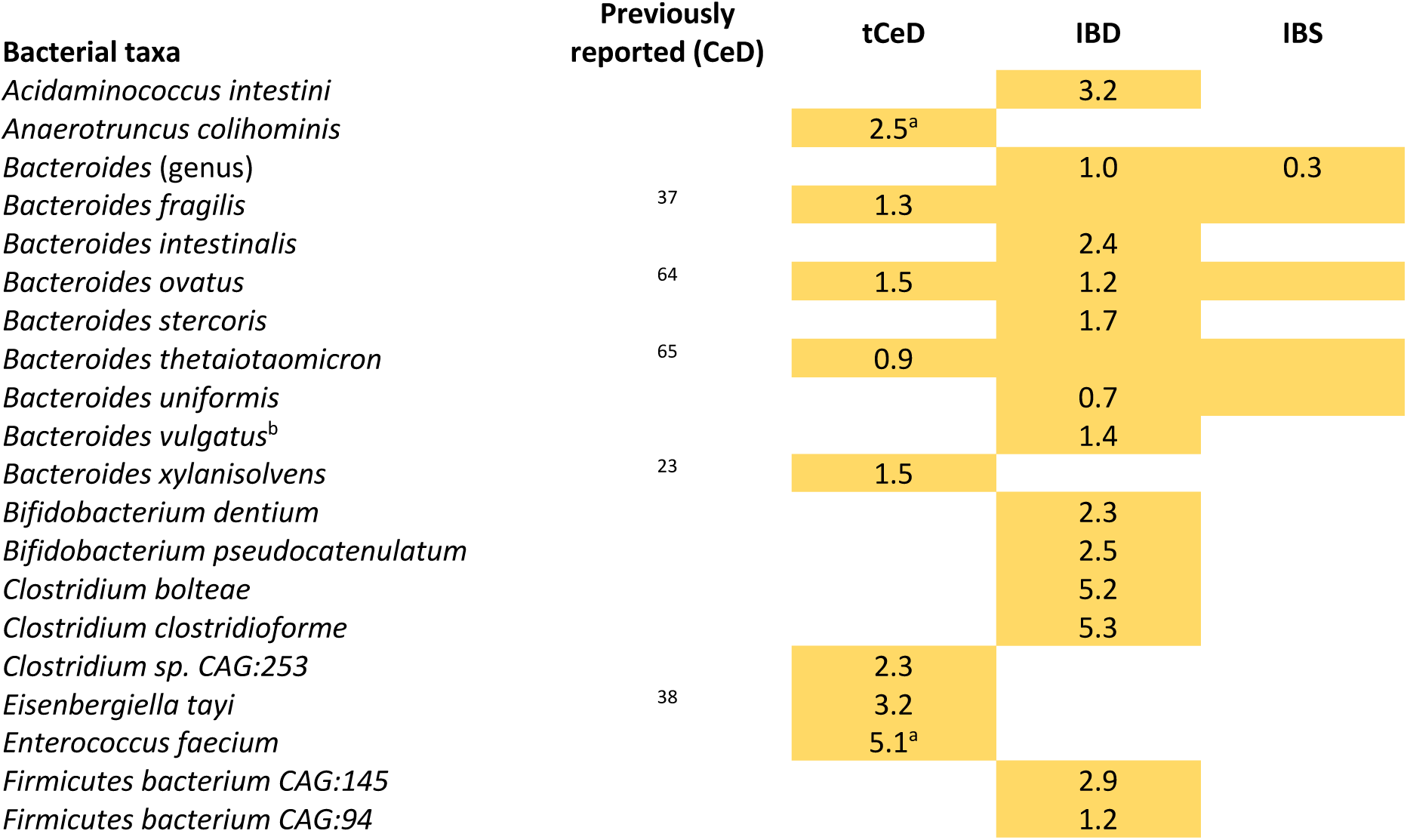
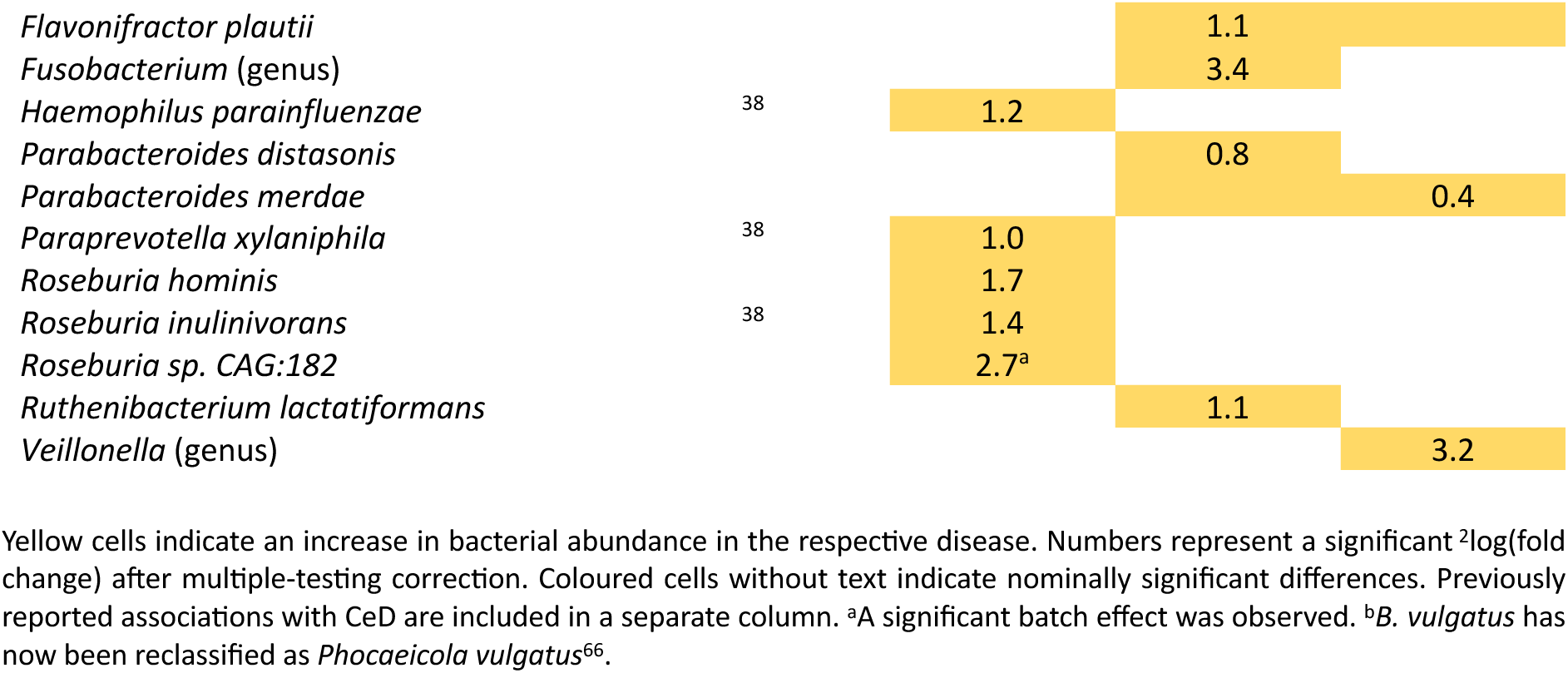
Taxa with significantly altered abundance in tCeD, IBD, and IBS.

**Table 3:**
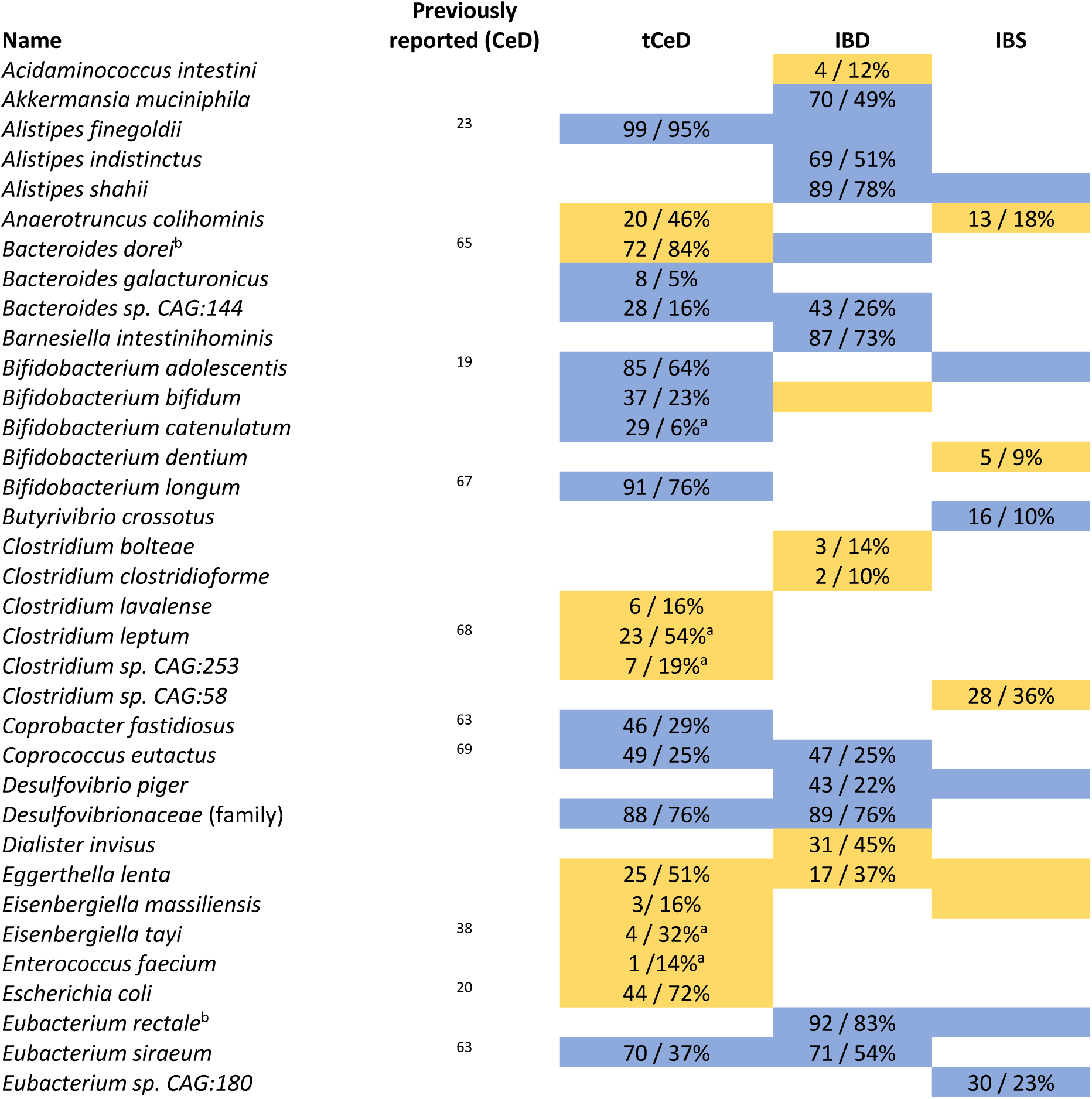
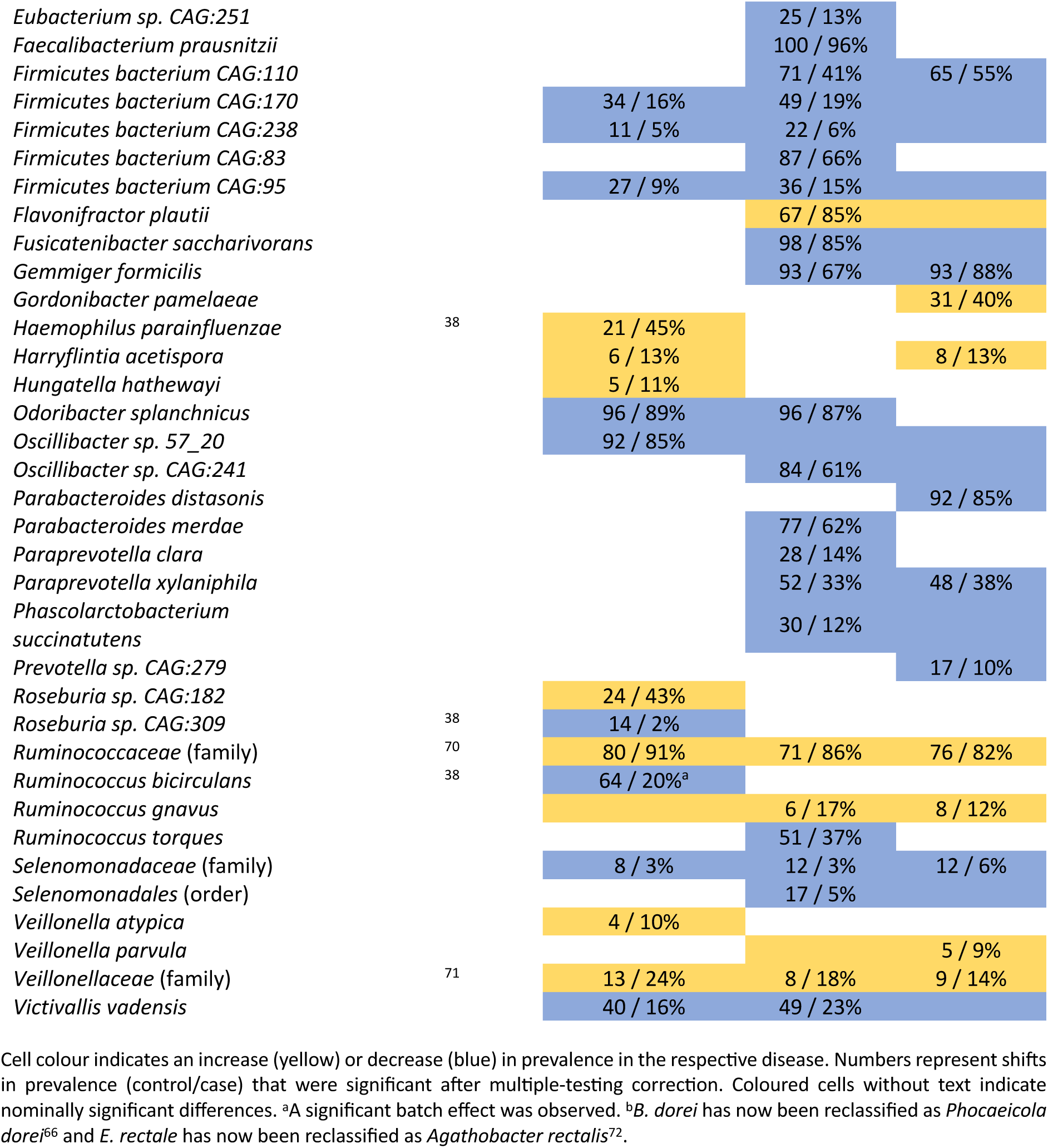
Taxa with significantly altered prevalence in tCeD, IBD, and IBS.

### Taxonomic analyses reveal both known and novel associations with tCeD

Our results confirm multiple previous reports of differences in bacterial abundance and/or prevalence between tCeD and controls (Table 2, Table 3), including enrichment of *H. parainfluenzae*^38^*, R. inulinivorans*^38^, *E. coli*^21,62^, and *Bacteroides* species^21,37,38^ and decreased prevalence of *R. bicirculans*^38^, *C. eutactus*^37^, *C. fastidiosus*^38,63^, *Roseburia sp. CAG:309*^38^, and several *Bifidobacterium* species^21,38,62^. We also identify several novel species, including an increase in *Clostridium sp. CAG:253*, *R. hominis*, and *E. lenta*.

### GFD duration partially explains the observed differences

Although we are primarily reporting on the microbiome composition of tCeD patients, we attempted to assess the contribution of a GFD to the increased abundance of the species shown in Table 2. After correction for covariates, the duration of GFD adherence positively correlated with increased abundances of *Paraprevotella xylaniphila* and *Roseburia sp. CAG:182* (Spearman rank correlation, *p_adj_* < 0.05; Figure 2). Moreover, given that the majority of the 13 species tested exhibit a positive correlation, these findings imply that a GFD does play a role in several of the observed species-level associations. To further confirm the effect of a GFD on the selected species, we compared 11 controls who reported adhering to a GFD to 55 matched controls who reported no specific diet. Among these controls, a GFD was also positively correlated with most of the studied species, including *P. xylaniphila* and *Roseburia sp. CAG:182* (Supp. Table S4). Notably, we were unable to replicate the adverse impact of a GFD on the abundance of *H. parainfluenzae* seen in other research^35,38^.

**Figure 2:**
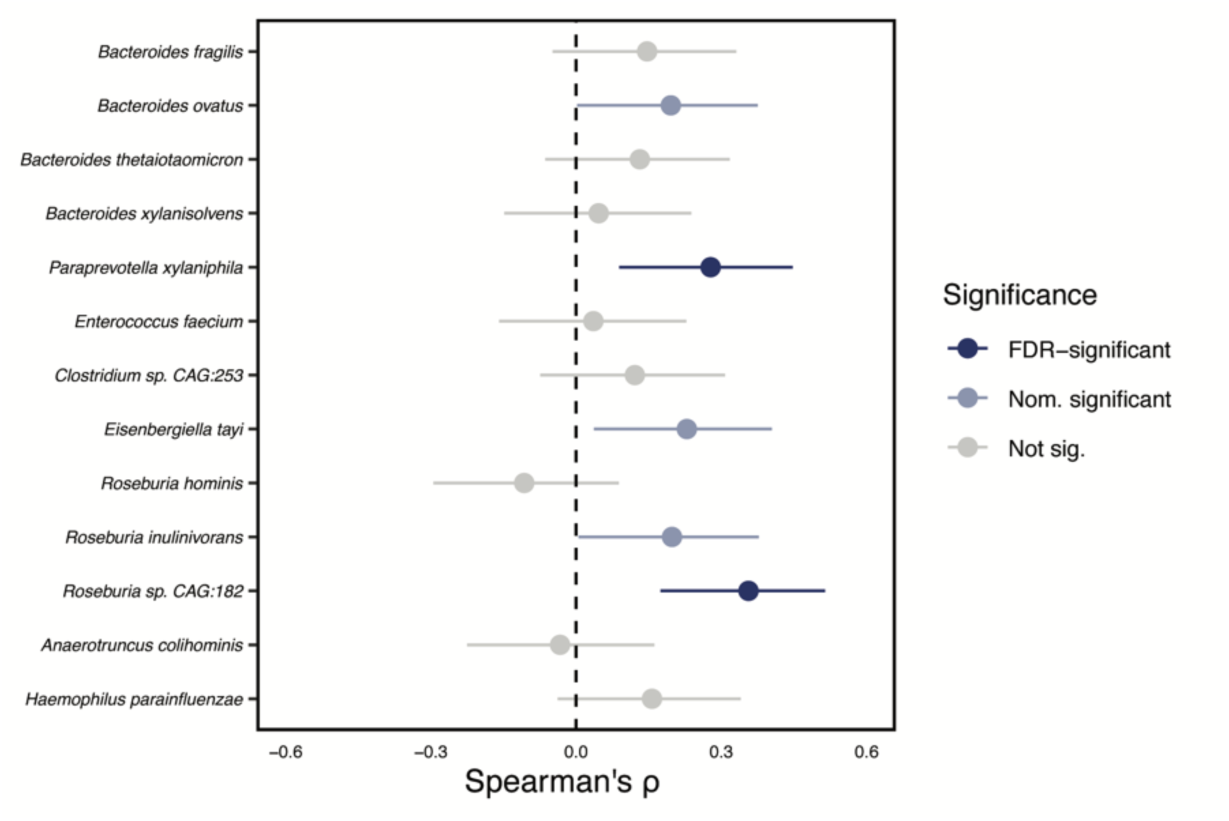
Spearman correlation coefficients between GFD duration and species abundance for all the species significantly enriched in tCeD patients. Nominal (p < 0.05) and corrected (p_adj_ < 0.05) significance are indicated in light and dark blue, respectively.

### Differences in metabolic pathways are largely driven by specific bacterial taxa

After establishing the taxonomic landscape in each of the subgroups analysed in this study, we assessed the functional potential of the associated microbiota by mapping the identified gene families to metabolic pathways in the MetaCyc database (Supp. Tables S5, S6, S7). Although we found several differences between tCeD patients and controls, very few were unique to tCeD and most showed considerable concordance with IBD and IBS (Table 4). These similarities can probably be explained by the fact that most of the differentially abundant metabolic pathways are strongly associated to differentially abundant taxa, with *Bacteroides* species and *E. coli* explaining most of the observed changes (Table 4, Supp. Figure S1, and Supp. Table S8). Furthermore, even the pathways with a lower abundance in tCeD compared to controls (Table 4, blue) appear to be primarily driven by an increase in bacteria from the *Bacteroidetes* phylum (now renamed *Bacteroidota*^17^) that replaces bacteria carrying this metabolic potential, reflecting the compositional nature of these data. Since pathway abundance seems to mainly reflect taxonomic changes rather than functional enrichment across several taxa, we do not consider it likely that these pathways are relevant in the context of the diseases studied.

**Table 4:**
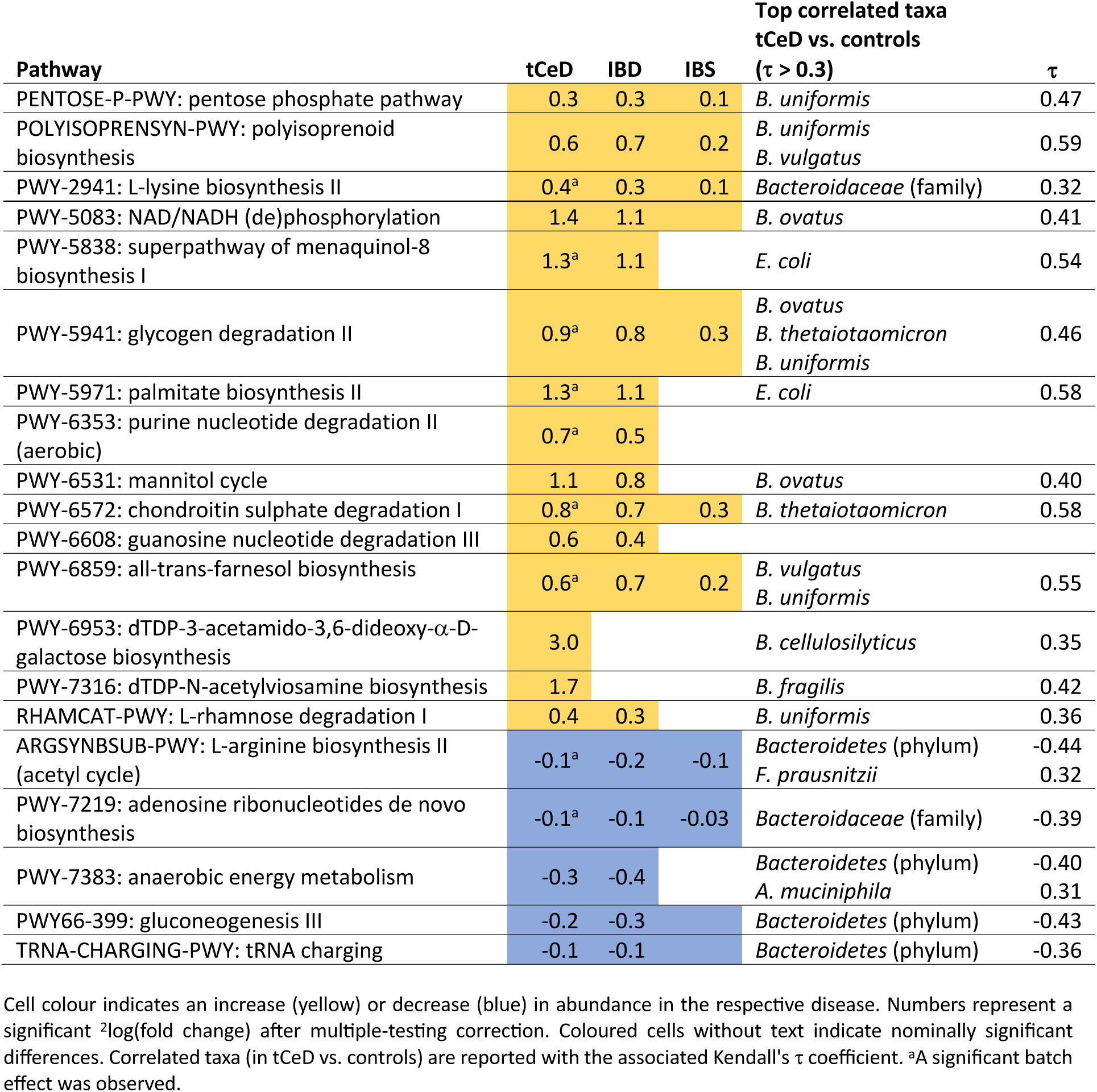
Metabolic pathways differentially abundant in tCeD.

### The tCeD faecal microbiome is less uniform than that of controls

Since we had identified several bacterial species specifically enriched in tCeD, we suspected that tCeD patients’ microbiomes would display a distinct CeD signature and be more uniform than those of non-CeD controls. Moreover, we also expected that the restricted GFD in tCeD could lead to a more homogeneous microbiome. To test this hypothesis, we first calculated

Bray-Curtis dissimilarity between samples, a measure of beta diversity, and performed principal coordinate analyses (Figure 3A). The principal coordinate plots for IBD and IBS are shown in Supp. Figure S2. Not surprisingly, the average overall composition of the tCeD microbiome differed significantly from that of controls (PERMANOVA, R^2^=0.003, *p* < 0.05). However, when we tested which group was more homogeneous, i.e. which group displayed the lowest median within-group dissimilarity (Figure 3B), we found that the microbiome composition of any two arbitrarily selected unrelated tCeD patients tended to be more dissimilar than that of two unrelated controls (permutation test, *p* < 0.05). Interestingly, we observed the same phenomenon, where patients have a less uniform microbiome structure than controls, among self-reported IBD patients (*p* < 0.05; Figure 3B). In contrast, the distributions of Bray-Curtis dissimilarity scores among controls and self-reported IBS patients were indistinguishable (Figure 3B).

**Figure 3:**
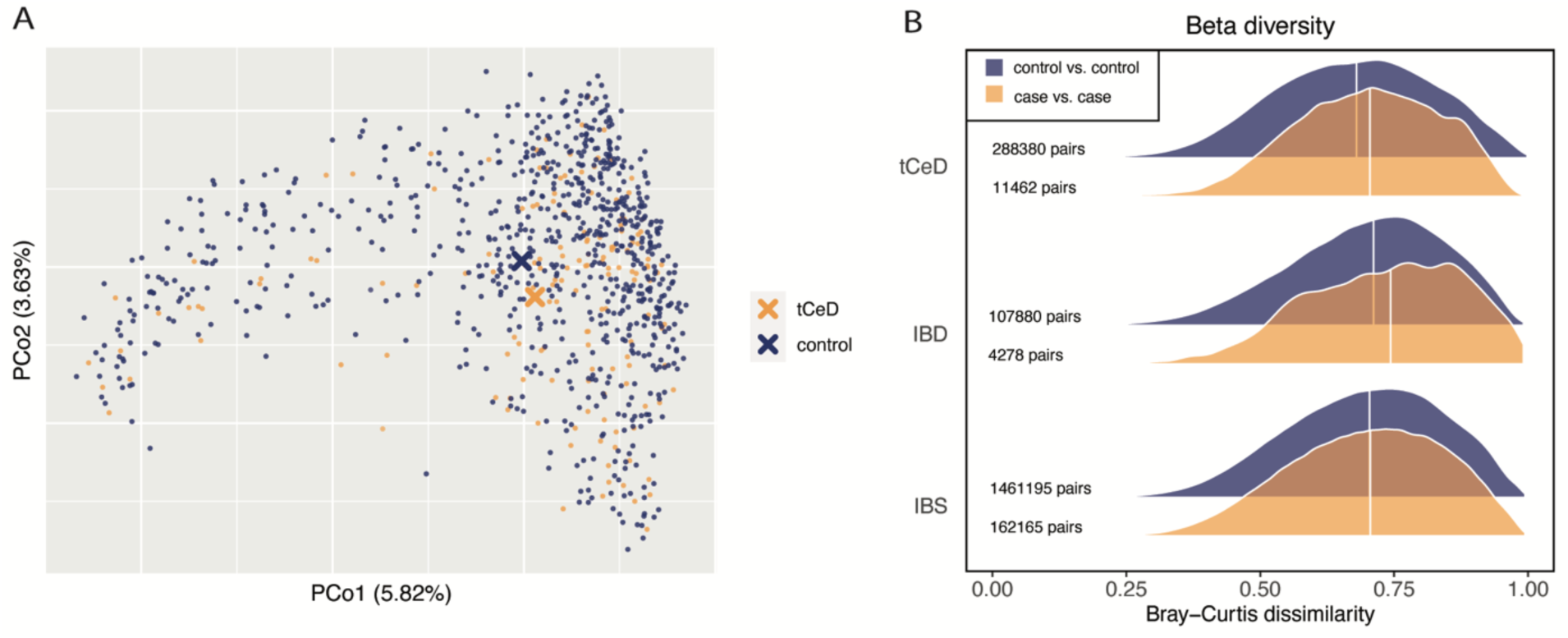
Beta diversity in tCeD vs. controls using Bray-Curtis dissimilarity as a distance metric. A. Principal coordinate plot of tCeD samples and matched controls. B. Within-group dissimilarity histograms for tCeD (top), IBD (middle), and IBS (bottom). Blue: pairs of unrelated controls. Orange: pairs of unrelated patients. Values at left are the number of different permutations of two samples that fall into each group.

### tCeD dissimilarity is also present on subspecies level

We next aimed to assess whether the increased diversity we observed among tCeD patients was only present on entire microbiome level, or whether this effect also extended to strain level. Using StrainPhlAn to estimate the pairwise evolutionary distance between bacterial strains of the same species from two different participants, we were able to analyse strain-level diversity for 67 bacterial species (Supp. Table S9). For nine species, the interindividual evolutionary strain distance among tCeD patients tended (nominal *p* < 0.05) to be higher than among controls, with only *Alistipes putredinis* showing a tendency toward more diversity among controls. After multiple-testing correction, the increased diversity among tCeD patients remained significant (Figure 4) for the *Bacteroides* species *B. ovatus, B. uniformis*, and *B. xylanisolvens* and for *Eubacterium rectale* (now reclassified as *Agathobacter rectalis*^72^ although its exact taxonomic classification is disputed^73–75^).

**Figure 4:**
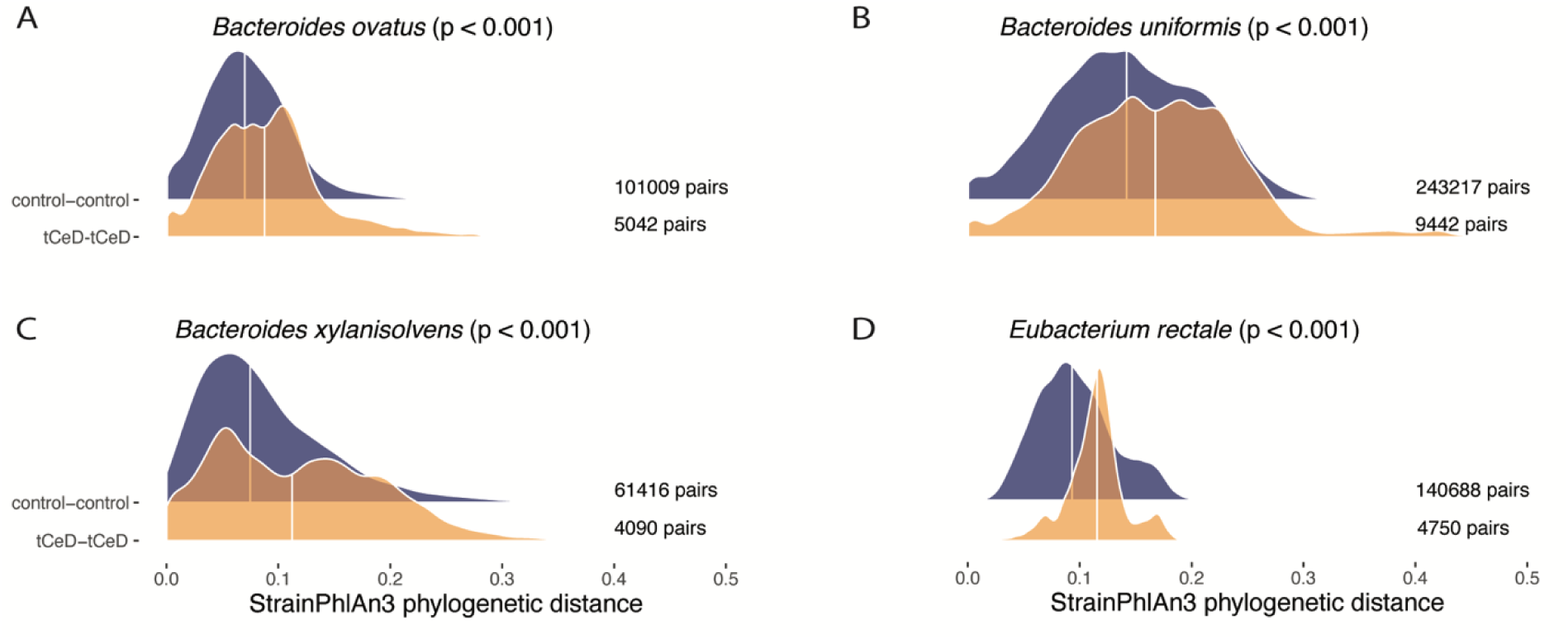
Distribution of pairwise genetic distances between strains of (A) *Bacteroides ovatus,* (B) *Bacteroides uniformis*, (C) *Bacteroides xylanisolvens*, and (D) *Eubacterium rectale*. Control vs. control (top) and tCeD vs. tCeD (bottom). Pairs of relatives were excluded from both comparisons. The significance of increased distance among tCeD patients is indicated by p-values from permutation tests (1000 iterations).

### European subspecies of *Eubacterium rectale* is reduced among tCeD patients

To further investigate within-species variation, we performed principal coordinate analyses, combined with PAM clustering. This approach revealed distinct subspecies clustering for 46 of the 67 species studied (Supp. Table S10). For *E. rectale*, the dominant cluster (Figure 5B, cluster 1) contained 61% of controls but only 33% of tCeD patients. Similarly, the dominant cluster in *Firmicutes bacterium CAG:83* (Supp. Figure S3, likely part of the *Oscillibacter* genus^76^) contained 69% of controls but 44% of tCeD patients (Fisher’s exact test, *p_adj_* < 0.05).

**Figure 5:**
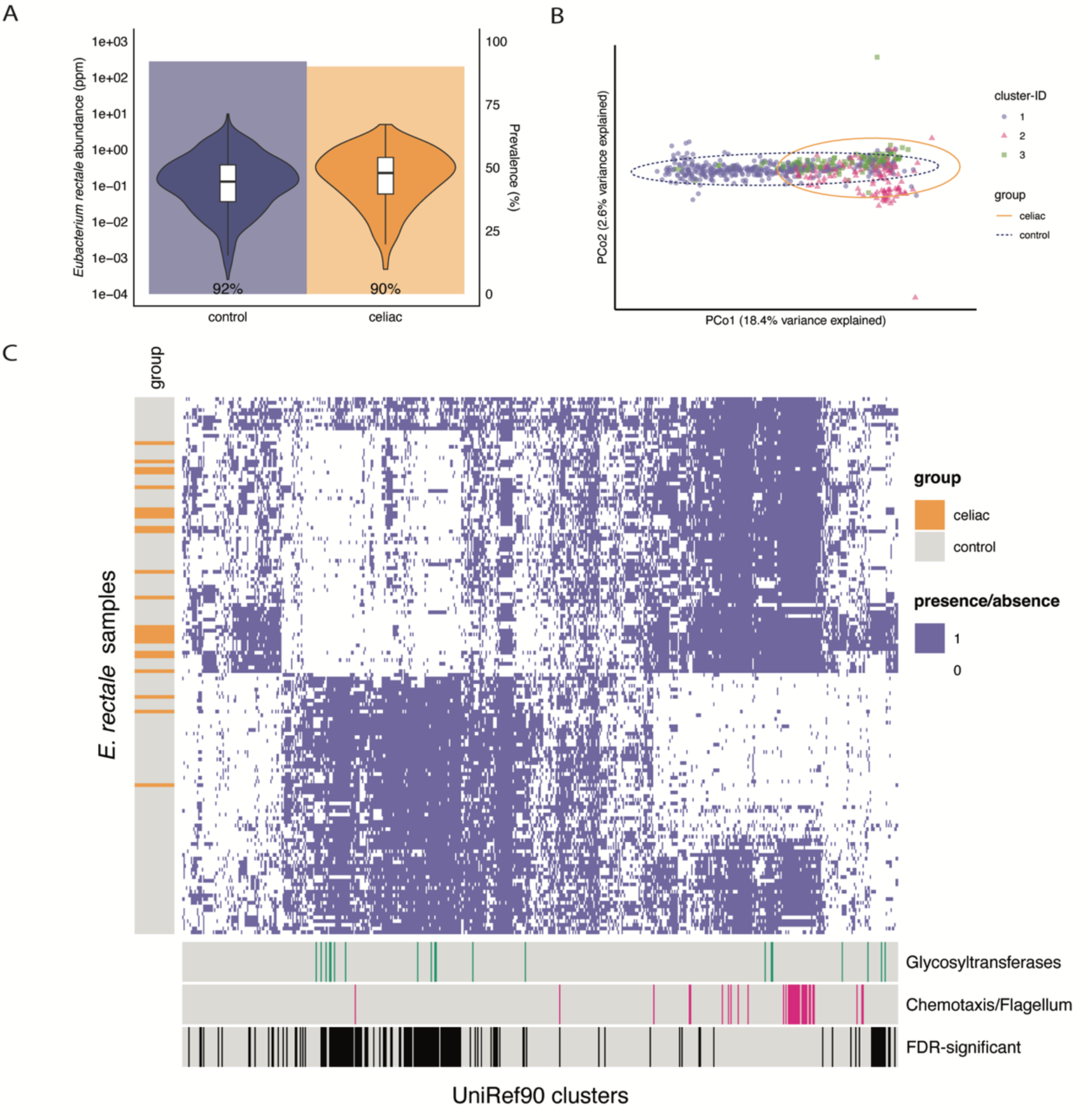
Strain-level differences of *Eubacterium rectale* between tCeD patients and controls. A. Abundance and prevalence of *E. rectale* in patients and controls. Violin plots represent the distribution of abundance (log-scale) in the two groups. Samples in which *E. rectale* was absent are excluded from the plot. Shaded bars in the background represent prevalence among all samples. B. Non-metric multidimensional scaling plots of pairwise genetic distances between *E. rectale* strains. Symbol shapes and colours indicate their assigned cluster (PAM clustering). Ellipses include 95% of the samples in the indicated group. C. Heatmap of the presence/absence of *E. rectale* genes in all the *E. rectale* strains analysed. Each row represents an *E. rectale* strain from one participant. CeD patients are indicated by the orange lines at left of the heatmap. Each column represents a UniRef90 cluster (gene) that is differentially prevalent in tCeD patients (nominal *p* < 0.05). Coloured bars below indicate whether the genes are predicted to encode glycosyltransferases (green) or flagellum-related proteins (pink). Black bars at bottom (FDR-significant) indicate the genes with *p_adj_* < 0.05.

Finally, we performed gene-level comparisons of cases and controls (PanPhlAn; Supp. Table S11), followed by enrichment analysis of protein family (Pfam) and gene ontology (GO) terms among nominally significant (*p* < 0.05) genes (Supp. Table S12). With this analysis, we aimed to identify gene-level explanations for the strain-level observations (increased interindividual differences and depletion of the dominant *E. rectale* cluster). The only class enriched among *E. rectale* genes that had a lower prevalence in tCeD was PF00535 (glycosyl transferase family 2), while over-prevalent genes were enriched for terms related to flagellum assembly and organisation, chemotaxis, protein targeting and secretion, and phosphoenolpyruvate-dependent sugar phosphotransferase systems (Figure 5C, Supp. Table S12). Karcher et al. previously described an *E. rectale* subspecies found almost exclusively in Europe (ErEurope) that is characterised by the presence of several genes from glycosyl transferase family 2 and the absence of motility operons, including chemotaxis and flagellum genes^77^. We therefore conclude that ErEurope is less prevalent in tCeD patients.

### Gene families related to transmembrane transport and DNA repair are less prevalent in multiple species in tCeD patients

In addition to the functional classes above, 139 other *E. rectale* genes were differentially prevalent in tCeD patients compared to controls (38 higher, 101 lower). Furthermore, genes from *Alistipes putredinis* (168 higher, 96 lower) and *Bifidobacterium adolescentis* (4 higher, 6 lower) were also found to be associated with tCeD (*p_adj_* < 0.05; Supp. Table S11). Interestingly, the *B. adolescentis* strains found in tCeD less frequently encode a LacI/GalR-family transcriptional regulator and galactose-1-phosphate uridyltransferase GalT, part of the Leloir pathway for galactose metabolism. This finding highlights another potential implication of the altered carbohydrate availability during inflammation and/or GFD^78^.

When we analysed strain-level differences between tCeD and controls in each of the seven species separately, no terms were enriched for any species beyond the glycosyltransferases and motility-related genes discussed above. To assess whether there were any bacterial gene or protein classes enriched across species, we performed the analysis after pooling the gene-level data of all seven bacteria. Among the under-prevalent genes in tCeD, we identified two new enriched classes. Firstly, 23 of 150 genes from GO:0055085 (transmembrane transport) were under-prevalent in tCeD. Many of the contributing genes, coming from six different species, were annotated to encode permeases, some with a predicted specificity for carbohydrates. We suggest that differences in nutrient availability during active CeD or GFD may play a role in the loss of these genes encoding carbohydrate-uptake proteins. Secondly, 9 of 21 genes encoding proteins from PF04002 (RadC-like JAB domain) were under-prevalent across all tested *Bacteroides* species and *Bacteroides vulgatus* (now reclassified as *Phocaeicola vulgatus*^66^). Proteins from this family have previously been speculated to have proteolytic and/or nuclease activity and to be involved in DNA repair^79^.

### Presence of DNA repair genes in four species results in lower mutational drift

Next, we assessed whether the reduced prevalence of DNA repair genes containing the RadC-like JAB domain in tCeD could be associated with increased interindividual strain variation. Indeed, the number of genes in a specific strain encoding a RadC-like JAB domain was negatively correlated (Figure 6) with the median distance of that strain to strains of the same species within the same sample group (tCeD and control). This negative correlation was observed in all species tested (*B. ovatus*, *B. uniformis*, *B. vulgatus*, and *B. xylanisolvens*) and significant in *B. uniformis* and *B. vulgatus,* the two species with the most data points (linear mixed-effects model, *p* < 0.05). Moreover, for all four species, there was a negative correlation in both the tCeD group and controls, in line with our hypothesis that these DNA repair proteins are limiting the mutation rate in these species.

**Figure 6:**
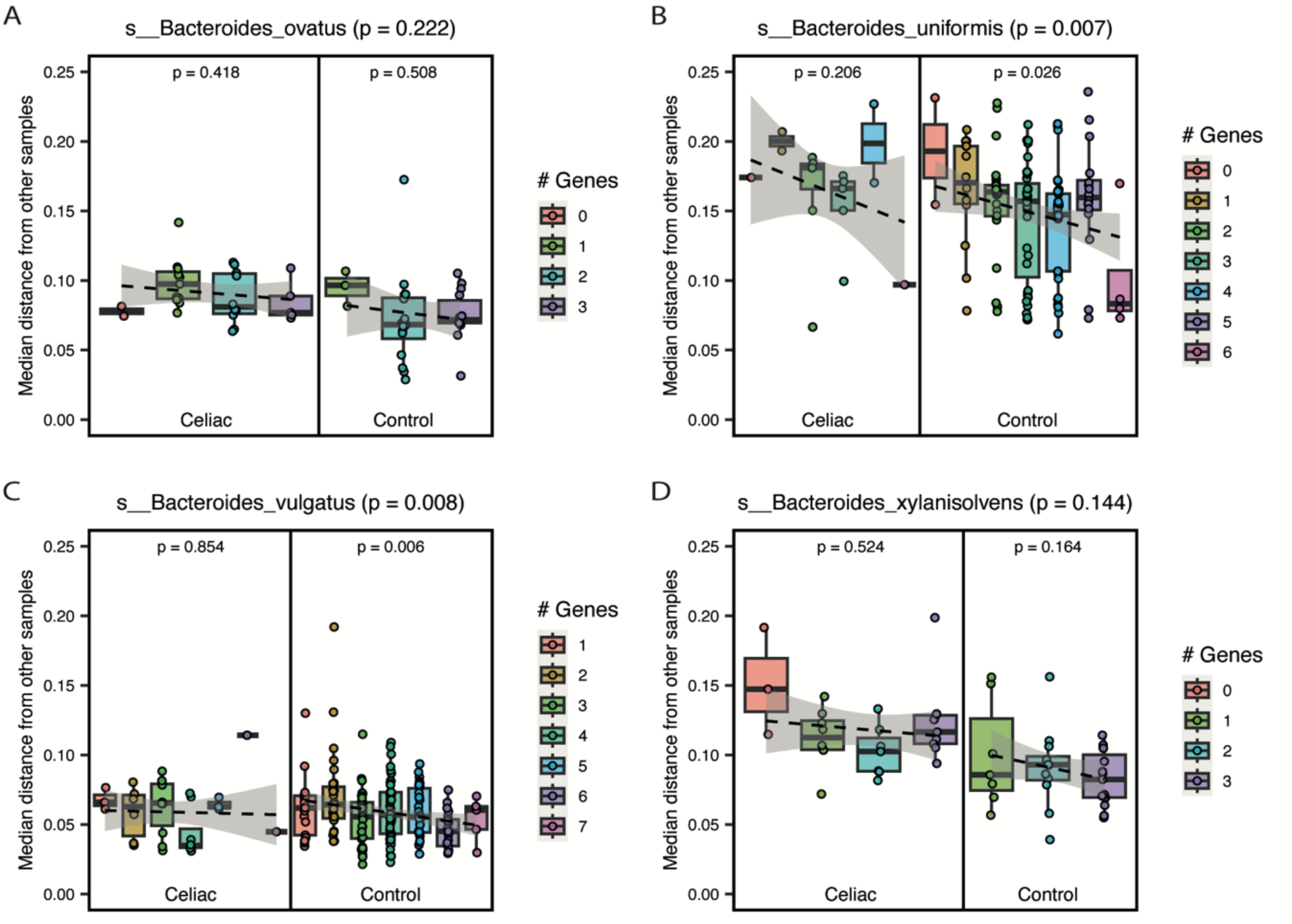
Correlation between prevalence in a specific sample of genes encoding the RadC-like JAB domain (’# Genes’) and the median mutational distance (StrainPhlAn) from other strains within the same sample group for (A) *Bacteroides ovatus*, (B) *Bacteroides uniformis*, (C) *Bacteroides vulgatus,* and (D) *Bacteroides xylanisolvens*. Significance within sample groups is indicated by the p-values inside the plot. The p-values in the plot title report the association in both groups combined (linear mixed-effects model).

## Discussion

We performed a comprehensive analysis of the largest cohort of metagenomics-sequenced CeD microbiota to date, comparing the faecal microbiota of 152 tCeD patients with 760 controls matched by age, sex, and sequencing depth. Our approach allows for microbiome classification at the strain level and provides insights into bacterial genes and pathways.

### Taxonomic and functional profiling

We aimed to compare taxonomic profiles at the species level with those of previous CeD microbiome studies. In summary, our data (Table 2 and Table 3) confirm previous findings in tCeD patients, demonstrating an increase in several *Bacteroides* species, *H. parainfluenzae, R. inulinivorans*, and *E. coli* and a decrease in various bifidobacteria, *R. bicirculans*, *C. eutactus,* and *C. fastidiosus*^21,37,38,62^. Despite a lower prevalence of the major butyrigenic bacterium *Butyrivibrio crossotus*^80^ (Table 3), we observed novel positive associations with other butyrate-producing bacteria including *Clostridium sp. CAG:253, R. hominis,* and *C. lavalense* in tCeD patients in our cohort. Butyrate, a short-chain fatty acid derived from the fermentation of dietary fibres, is an important modulator of immune and barrier functions, which are relevant for CeD^80–82^. Higher production of butyrate by the CeD microbiome aligns with observations by Baldi et al. where CeD patients displayed slightly higher circulating butyrate fractions^83^. However, Tjellström et al. showed that one year of GFD adherence restored butyrate levels to normal in paediatric CeD patients^84^. We speculate that the reducing effect of a GFD on butyrate production may not necessarily reflect restoration of the microbiome to a pre-CeD state. It may instead be mediated by other butyrigenic species so that the effects of disease and diet largely cancel each other out. Changes in butyrate producers may be a result of the altered fibre content of a GFD^85–87^.

Further taxonomic analyses revealed that tCeD, IBD, and IBS were all characterised by an increase in the *Bacteroides* genus, which drives the remarkably similar changes in metabolic pathways observed in each disease group. Moreover, it seems that all (moderately) under-abundant pathways can be explained by this (moderate) increase of bacteria from the *Bacteroidetes* phylum, suggesting that this is likely an artefact caused by the compositional nature of the data (Table 4, Supp. Figure S1).

Besides genus *Bacteroides*, however, the different disease groups studied each show highly distinct microbiome profiles (Table 2, Table 3, Figure 1), indicating that the general inflammatory milieu and increased barrier permeability associated with both IBD and CeD do not result in similar microbiome profiles. Although we show that the GFD contributes to an altered faecal microbiome (Figure 2), the current data does not allow us to separate the consequences of CeD and/or GFD from characteristics of the microbiome that contribute to the onset of CeD.

Although most metabolic pathways appear to be altered because the bacteria encoding them are altered in tCeD, we cannot dismiss the possibility that certain bacteria are enriched because they encode those metabolic pathways. Indeed, other studies have reported several *Bacteroides* species to be enriched in active^21,57,88^ and treated CeD patients^21,37,57^ as well as at-risk individuals^23,89,90^. In these studies, *B. fragilis* was speculated to be involved in pathogenesis^64^. Although these studies did not perform multiple-testing correction, taken together they do suggest that the overabundance of *Bacteroides* species is not solely caused by GFD and that some of the pathways we identified might therefore be relevant in CeD pathogenesis. Additionally, the 1.3-fold enrichment of genes involved in the rhamnose degradation pathway we observe (Table 4) could be related to differences in carbohydrate supply from the host diet. Furthermore, we found two pathways uniquely affected in tCeD, dTDP-N-acetylviosamine biosynthesis (3.3-fold enriched in tCeD) and dTDP-3-acetamido-3,6-dideoxy-α-D-galactose biosynthesis pathways (7.8-fold enriched in tCeD), which also represent the two largest functional differences compared to controls. Both these pathways use α-D-glucopyranose 1-phosphate, a product of glycogen degradation, as a substrate. Since both these pathways and, as in IBD, glycogen degradation pathways are overabundant in tCeD and associated with various *Bacteroides* species, it could be that the ability to survive in harsh conditions with low carbohydrate availability gives these bacteria a competitive advantage over other bacteria^91^. Only two pathways enriched in tCeD do not appear to be driven by specific bacterial taxa, PWY-6353 and PWY-6608 (Table 4), both related to purine nucleotide degradation.

### Interindividual variation

Besides the species-specific differences between tCeD patients and controls, we found an increased interindividual diversity among patients compared to controls (Figure 3). Thus, while we had expected that patients would share a specific microbiome signature related to either the disease or GFD, the inverse appears to be true. The tCeD microbiome deviates more from the average non-CeD microbiome (not shown) than that of controls, but these changes apparently occur in different directions between patients. Although we prefer not to use the term *dysbiosis*^92^, this observation could reflect the loss of certain healthy components of the microbiome that are compensated for differently in each individual. On the other hand, it could be that a gluten-*containing* diet, rather than disease or GFD, induces a microbiome signature. While the possibility of a loss of specific ‘healthy’ bacteria characterising CeD seems incompatible with the fact that we exclusively observed overabundant bacteria in tCeD, presence-absence analyses did reveal some bacteria with reduced prevalence in tCeD, such as several species of bifidobacteria (generally regarded as ‘healthy’ bacteria^93^) and *R. bicirculans*, *E. siraeum*, and *C. eutactus* (Table 3).

Interestingly, the increased diversity we observe in patients is also visible in the strain-level analyses, where, for several bacteria, the average evolutionary distance between strains of the same species in two unrelated tCeD patients was larger than that between strains in two control individuals (Supp. Table S9). This phenomenon was observed most clearly (p_adj_ < 0.05) in *Eubacterium rectale*, *Bacteroides ovatus*, *Bacteroides uniformis*, and *Bacteroides xylanisolvens* (Figure 4). But it also appears to be more general, as 48 of the 67 species analysed (72%) show the same trend. The most plausible explanation we see is that the inflammation and oxidative stress accompanying CeD and gluten intake^94^ lead to a higher mutational burden in the gut microbiome. In line with this hypothesis, transcriptome analysis of intestinal epithelial cells recently revealed the upregulation of host pathways related to DNA double-strand break repair in active CeD compared to controls^95^. Moreover, we find *Bacteroides* strains (multiple species) that carry the RadC-like JAB domain involved in DNA repair^79^ to be more prevalent in controls than in tCeD (Supp. Tables S11 and S12) and that the presence of these repair proteins in *Bacteroides* strains limits their mutation rate and thereby inter-strain variation (Figure 6). These observations together suggest that under stressful conditions it may be beneficial to these bacteria to increase the genetic diversity of their offspring. Notably, we observe the same trend in the self-reported IBD samples, potentially reflecting the mutational burden of an inflamed environment. Indeed, it is an established strategy among bacteria to upregulate their evolution rate in response to stress^96^. Examples of this strategy are the widespread SOS-response, utilising an error-prone DNA polymerase^97^, and the activation of competence for transformation during DNA replication stress^98,99^. It is unclear whether the increase in mutational load we observe in the tCeD microbiota is an indication of residual inflammation or merely a result of earlier periods of inflammation.

Intriguingly, while *Eubacterium rectale* also shows clear evidence of (seemingly) random mutagenesis (Figure 4D, Supp. Figure S4), we also observed that the ErEurope subspecies^77^ was almost absent in tCeD, while nearly half of the controls carried this subspecies. Importantly, ErEurope lacks many motility genes and has a distinct carbohydrate metabolism potential. Karcher et al. have suggested that ErEurope utilises complex plant-derived carbohydrates more efficiently^77^. As no interaction with diet was previously observed for ErEurope, most of the different *E. rectale* strains do not co-colonise^77^, and we observe no differences in prevalence or abundance between tCeD patients and controls (Figure 5A), it is possible that the strain-level difference in *E. rectale* may exist prior to GFD. It remains unclear whether the lack of motility limits ErEurope’s ability to access carbohydrates during inflammation, whether their carbohydrate metabolism potential plays a role in gluten processing, or whether pre-existing genetic or environmental risk factors for CeD also affect *E. rectale* sub-strain colonisation success. Alternatively, as metabolites produced by certain *E. rectale* strains have been implicated in alleviating intestinal inflammation^100^, selection pressure may also be present. Finally, the overrepresentation of flagellated *E. rectale* strains in tCeD, combined with the reported enrichment of anti-flagellar antibodies in IBD patients^101^, could imply a role for flagellins in disease aetiology or manifestation, e.g. through activation of Toll-like receptor 5 and initiation of Th1 T cell responses^102^.

### Limitations

Since the CeD patients in our study strictly followed a GFD, our findings do not allow us to definitively ascertain the role of the microbiome in CeD onset or progression. This challenge is common in much of the existing research, where data interpretation is hindered by diet and the possibility that observed differences stem from studying patients with active CeD, potentially influenced by associated intestinal inflammation. Furthermore, our study is limited to faecal samples, which may not accurately reflect conditions in the small intestine^26^.

Despite these limitations, the current findings are a starting point for future work addressing which differences truly reflect changes in the small intestine, which differences precede disease onset, and what mechanisms are responsible for these differences in microbial composition. Additional future data, ideally obtained from prospective and longitudinal studies − including those on active CeD patients, healthy individuals adhering to a GFD, and/or at-risk individuals − will help tease apart the various influences of genetics, diet, and disease on the gut microbiome of CeD patients. A longitudinal study published in 2021 where data was available before CeD onset reported several differences, but this study had limited taxonomic resolution (16S sequencing) and statistical power (only 10 cases and 10 controls)^23^. Furthermore, as the significance levels in this earlier study appear not to have been corrected for multiple testing, it is difficult to assess whether the reported differences are beyond what would be expected by chance. However, although the microbiome analysis in the longitudinal, prospective TEDDY study^103^ also suffered from small sample size when they reported on it in 2018^24^, we expect that future updated analyses of this large CeD patient cohort^104,105^ will prove a valuable addition to our current findings.

### Conclusion

We characterised the microbiota of tCeD patients compared to matched controls, leading to confirmation of previously reported CeD–microbiome associations and identification of novel associations that may be linked to CeD, GFD, or both. Strain- and gene-level comparisons revealed a higher mutation frequency in several species, possibly induced by inflammation and/or oxidative stress. Data on the prevalence of the RadC-like JAB domain in multiple *Bacteroides* species suggest that increased mutational load may increase bacterial adaptability under stressful conditions. Finally, we found that the European subspecies of *Eubacterium rectale* is virtually absent in tCeD patients, a finding that may help shed light on some of the mechanisms that shape the microbiome in health and disease.

## Data availability

### Lifelines Dutch Microbiome Project (LL-DMP)

The raw microbiome sequencing data, processed microbiome data (including taxonomy and pathways), and basic phenotypes (including age, sex and BMI) from LL-DMP are available at the European Genome-Phenome Archive (EGA) under accession EGAS00001005027. Access to these dataset can be requested via the page of the EGA data access committee EGAC00001001996. The phenotype data can be requested, for a fee, by filling the application form at https://www.lifelines.nl/researcher/how-to-apply/apply-here. For questions about accessing other Lifelines data, researchers can contact Lifelines at. For questions regarding the selected LL-DMP samples for this study, please contact the corresponding author of this study (S.W.).

### Celiac Disease Northern Netherlands (CeDNN)

Access to the raw microbiome sequencing data, processed microbiome data (including taxonomy and pathways), and basic phenotypes (including age, sex and BMI) from CeDNN are available at the EGA under accession [SUBMISSION PENDING]. The phenotype data can be requested by contacting the corresponding author of this study (S.W.).

## Code availability

Open source codes and scripts used for the processing of raw metagenomic sequencing data are available at the GitHub repository https://github.com/GRONINGEN-MICROBIOME-CENTRE/GMH_MGS_pipeline. Only parameters referring to *bioBakery 3* tools (*MetaPhlAn3, HUMAnn3, StrainPhlAn3* and *PanPhlAn3*) apply for the current study. To facilitate the re-use of the codes, the repository also includes example datasets that enable users to test the codes without the need to apply for access to phenotypes.

## Acknowledgements

We would like to thank the participants of CeDNN and Lifelines for making this research possible. We also acknowledge the services of Lifelines Cohort Study and the contributing research centres delivering data to Lifelines. We thank the Center for Information Technology of the University of Groningen (RUG) and the Genomics Coordination Center (UMCG and RUG) for their support and for providing access to their respective high-performance computing clusters. We also thank the MOLGENIS team for data management and analysis support. We thank Kate Mc Intyre for editing the manuscript.

This work was supported by the Netherlands Organization for Scientific Research (NWO) under Grants 024.003.001 (Netherlands Organ-on-Chip Initiative; J.S., H.S., C.W., J.F., S.W.), 024.004.017 (Exposome-NL; A.Z.), VI.C.202.022 (VICI; J.F.), 016.178.056 (VIDI; A.Z.), 016.136.308 (VIDI; R.K.W.), and 016.171.047 (VIDI; I.H.J.); the European Research Council (ERC) under Grants 101001678 (Consolidator; J.F.) and 715772 (Starting; A.Z.); the EU Horizon Europe under Grant INITIALISE (A.Z.); the Dutch Digestive Foundation under Grant MLDS D16–14 (Diagnostics, R.K.W.); the Dutch Heart Foundation under Grant CVON2018-27 (IN-CONTROL; A.Z., J.F.); the Natural Science Foundation of China under Grant 32270077 (L.C.); the Natural Science Foundation of Jiangsu under Grant BK20220709 (L.C.); the Nanjing Medical University under Grant BK20220709 (Starting; L.C.) and Stichting Ammodo (AMMODO Science Award 2023; J.F.). In addition, I.T. was supported by a MD/PhD scholarship from the Junior Scientific Masterclass (Graduate School of Medical Sciences, University Medical Center Groningen, and University of Groningen) and a KNAW Ter Meulen grant; L.C. was supported by the De Cock-Hadders Foundation and I.H.J. is supported by a Rosalind Franklin Fellowship from the University of Groningen.

## References

1. Ludvigsson JF, Lefler DA, Bai JC, Biagi F, Fasano A, Green PHR, Hadjivassiliou M, Kaukinen K, Kelly CP, Leonard JN, et al. The Oslo definitions for coeliac disease and related terms. Gut 2013; 62:43–52.

2. Lebwohl B, Sanders DS, Green PHR. Coeliac disease. Lancet 2018; 391:70–81.

3. Lebwohl B, Ludvigsson JF, Green PHR. Celiac disease and non-celiac gluten sensitivity. BMJ 2015; 351:h4347.

4. Gujral N, Freeman HJ, Thomson ABR. Celiac disease: prevalence, diagnosis, pathogenesis and treatment. World J Gastroenterol 2012; 18:6036–59.

5. Rewers M. Epidemiology of celiac disease: what are the prevalence, incidence, and progression of celiac disease? Gastroenterology 2005; 128:S47–51.

6. van de Wal Y, Kooy Y, van Veelen P, Peña S, Mearin L, Papadopoulos G, Koning F. Selective deamidation by tissue transglutaminase strongly enhances gliadin-specific T cell reactivity. J Immunol 1998; 161:1585–8.

7. Meling MT, Kleppa L, Besser HA, Khosla C, du Pré MF, Sollid LM. Enterocyte-derived and catalytically active transglutaminase 2 in the gut lumen of mice: implications for celiac disease. Gastroenterology 2024; 167:1026–1028.e4.

8. Kårhus LL, Thuesen BH, Skaaby T, Rumessen JJ, Linneberg A. The distribution of HLA DQ2 and DQ8 haplotypes and their association with health indicators in a general Danish population. United European Gastroenterol J 2018; 6:866–78.

9. Bodd M, Ráki M, Tollefsen S, Fallang LE, Bergseng E, Lundin KEA, Sollid LM. HLA-DQ2-restricted gluten-reactive T cells produce IL-21 but not IL-17 or IL-22. Mucosal Immunol 2010; 3:594–601.

10. Di Sabatino A, Vanoli A, Giuffrida P, Luinetti O, Solcia E, Corazza GR. The function of tissue transglutaminase in celiac disease. Autoimmun Rev 2012; 11:746–53.

11. Troncone R, Jabri B. Coeliac disease and gluten sensitivity. J Intern Med 2011; 269:582–90.

12. Al-Toma A, Volta U, Auricchio R, Castillejo G, Sanders DS, Cellier C, Mulder CJ, Lundin KEA. European Society for the Study of Coeliac Disease (ESsCD) guideline for coeliac disease and other gluten-related disorders. United European Gastroenterol J 2019; 7:583–613.

13. Kumar V, Wijmenga C, Withoff S. From genome-wide association studies to disease mechanisms: celiac disease as a model for autoimmune diseases. Semin Immunopathol 2012; 34:567–80.

14. van Heel DA, Franke L, Hunt KA, Gwilliam R, Zhernakova A, Inouye M, Wapenaar MC, Barnardo MCNM, Bethel G, Holmes GKT, et al. A genome-wide association study for celiac disease identifies risk variants in the region harboring IL2 and IL21. Nat Genet 2007; 39:827–9.

15. Hunt KA, Zhernakova A, Turner G, Heap GAR, Franke L, Bruinenberg M, Romanos J, Dinesen LC, Ryan AW, Panesar D, et al. Newly identified genetic risk variants for celiac disease related to the immune response. Nat Genet 2008; 40:395–402.

16. Dubois PCA, Trynka G, Franke L, Hunt KA, Romanos J, Curtotti A, Zhernakova A, Heap GAR, Adány R, Aromaa A, et al. Multiple common variants for celiac disease influencing immune gene expression. Nat Genet 2010; 42:295–302.

17. Oren A, Garrity GM. Valid publication of the names of forty-two phyla of prokaryotes. Int J Syst Evol Microbiol 2021; 71.

18. Collado MC, Calabuig M, Sanz Y. Differences between the fecal microbiota of coeliac infants and healthy controls. Curr Issues Intest Microbiol 2007; 8:9–14.

19. Sanz Y, Sánchez E, Marzotto M, Calabuig M, Torriani S, Dellaglio F. Differences in faecal bacterial communities in coeliac and healthy children as detected by PCR and denaturing gradient gel electrophoresis. FEMS Immunology & Medical Microbiology 2007; 51:562–8.

20. Nadal I, Donant E, Ribes-Koninckx C, Calabuig M, Sanz Y. Imbalance in the composition of the duodenal microbiota of children with coeliac disease. J Med Microbiol 2007; 56:1669–74.

21. Collado MC, Donat E, Ribes-Koninckx C, Calabuig M, Sanz Y. Specific duodenal and faecal bacterial groups associated with paediatric coeliac disease. Journal of Clinical Pathology 2009; 62:264–9.

22. Collado MC, Donat E, Ribes-Koninckx C, Calabuig M, Sanz Y. Imbalances in faecal and duodenal *Bifidobacterium* species composition in active and non-active coeliac disease. BMC Microbiol 2008; 8:232.

23. Leonard MM, Valitutti F, Karathia H, Pujolassos M, Kenyon V, Fanelli B, Troisi J, Subramanian P, Camhi S, Colucci A, et al. Microbiome signatures of progression toward celiac disease onset in at-risk children in a longitudinal prospective cohort study. Proceedings of the National Academy of Sciences 2021; 118:e2020322118.

24. Stewart CJ, Ajami NJ, O’Brien JL, Hutchinson DS, Smith DP, Wong MC, Ross MC, Lloyd RE, Doddapaneni H, Metcalf GA, et al. Temporal development of the gut microbiome in early childhood from the TEDDY study. Nature 2018; 562:583–8.

25. Caminero A, McCarville JL, Galipeau HJ, Deraison C, Bernier SP, Constante M, Rolland C, Meisel M, Murray JA, Yu XB, et al. Duodenal bacterial proteolytic activity determines sensitivity to dietary antigen through protease-activated receptor-2. Nat Commun 2019; 10:1198.

26. Constante M, Libertucci J, Galipeau HJ, Szamosi JC, Rueda G, Miranda PM, Pinto-Sanchez MI, Southward CM, Rossi L, Fontes ME, et al. Biogeographic variation and functional pathways of the gut microbiota in celiac disease. Gastroenterology 2022; 163:1351–1363.e15.

27. Caminero A, Galipeau HJ, McCarville JL, Johnston CW, Bernier SP, Russell AK, Jury J, Herran AR, Casqueiro J, Tye-Din JA, et al. Duodenal bacteria from patients with celiac disease and healthy subjects distinctly affect gluten breakdown and immunogenicity. Gastroenterology 2016; 151:670–83.

28. Cardoso-Silva D, Delbue D, Itzlinger A, Moerkens R, Withoff S, Branchi F, Schumann M. Intestinal barrier function in gluten-related disorders. Nutrients 2019; 11:2325.

29. Matar A, Damianos JA, Jencks KJ, Camilleri M. Intestinal barrier impairment, preservation, and repair: an update. Nutrients 2024; 16:3494.

30. Wu S, Hashimoto-Hill S, Woo V, Eshleman EM, Whitt J, Engleman L, Karns R, Denson LA, Haslam DB, Alenghat T. Microbiota-derived metabolite promotes HDAC3 activity in the gut. Nature 2020; 586:108–12.

31. Petersen J, Ciacchi L, Tran MT, Loh KL, Kooy-Winkelaar Y, Croft NP, Hardy MY, Chen Z, McCluskey J, Anderson RP, et al. T cell receptor cross-reactivity between gliadin and bacterial peptides in celiac disease. Nat Struct Mol Biol 2020; 27:49–61.

32. Halfvarson J, Brislawn CJ, Lamendella R, Vázquez-Baeza Y, Walters WA, Bramer LM, D’Amato M, Bonfiglio F, McDonald D, Gonzalez A, et al. Dynamics of the human gut microbiome in inflammatory bowel disease. Nat Microbiol 2017; 2:1–7.

33. Zeng MY, Inohara N, Nuñez G. Mechanisms of inflammation-driven bacterial dysbiosis in the gut. Mucosal Immunol 2017; 10:18–26.

34. Sellitto M, Bai G, Serena G, Fricke WF, Sturgeon C, Gajer P, White JR, Koenig SSK, Sakamoto J, Boothe D, et al. Proof of concept of microbiome-metabolome analysis and delayed gluten exposure on celiac disease autoimmunity in genetically at-risk infants. PLoS One 2012; 7:e33387.

35. Sample D, Fouhse J, King S, Huynh HQ, Dieleman LA, Willing BP, Turner J. Baseline fecal microbiota in pediatric patients with celiac disease is similar to controls but dissimilar after 1 year on the gluten-free diet. JPGN Rep 2021; 2:e127.

36. Olivares M, Walker AW, Capilla A, Benítez-Páez A, Palau F, Parkhill J, Castillejo G, Sanz Y. Gut microbiota trajectory in early life may predict development of celiac disease. Microbiome 2018; 6:36.

37. Palmieri O, Castellana S, Bevilacqua A, Latiano A, Latiano T, Panza A, Fontana R, Ippolito AM, Biscaglia G, Gentile A, et al. Adherence to gluten-free diet restores alpha diversity in celiac people but the microbiome composition is different to healthy people. Nutrients 2022; 14:2452.

38. Francavilla A, Ferrero G, Pardini B, Tarallo S, Zanatto L, Caviglia GP, Sieri S, Grioni S, Francescato G, Stalla F, et al. Gluten-free diet affects fecal small non-coding RNA profiles and microbiome composition in celiac disease supporting a host-gut microbiota crosstalk. Gut Microbes 2023; 15:2172955.

39. Brouwer-Brolsma EM, Streppel MT, van Lee L, Geelen A, Sluik D, van de Wiel AM, de Vries JHM, van ‘t Veer P, Feskens EJM. A National Dietary Assessment Reference Database (NDARD) for the Dutch population: rationale behind the design. Nutrients 2017; 9:1136.

40. Gacesa R, Kurilshikov A, Vich Vila A, Sinha T, Klaassen M a. Y, Bolte LA, Andreu-Sánchez S, Chen L, Collij V, Hu S, et al. Environmental factors shaping the gut microbiome in a Dutch population. Nature 2022; 604:732–9.

41. Ho D, Imai K, King G, Stuart EA. MatchIt: nonparametric preprocessing for parametric causal inference. Journal of Statistical Software 2011; 42:1–28.

42. Imhann F, Van der Velde KJ, Barbieri R, Alberts R, Voskuil MD, Vich Vila A, Collij V, Spekhorst LM, Van der Sloot KWJ, Peters V, et al. The 1000IBD project: multi-omics data of 1000 inflammatory bowel disease patients; data release 1. BMC Gastroenterol 2019; 19:5.

43. Tigchelaar EF, Zhernakova A, Dekens JAM, Hermes G, Baranska A, Mujagic Z, Swertz MA, Muñoz AM, Deelen P, Cénit MC, et al. Cohort profile: LifeLines DEEP, a prospective, general population cohort study in the northern Netherlands: study design and baseline characteristics. BMJ Open 2015; 5:e006772.

44. McIver LJ, Abu-Ali G, Franzosa EA, Schwager R, Morgan XC, Waldron L, Segata N, Huttenhower C. bioBakery: a meta’omic analysis environment. Bioinformatics 2018; 34:1235–7.

45. Langmead B, Salzberg SL. Fast gapped-read alignment with Bowtie 2. Nat Methods 2012; 9:357–9.

46. Beghini F, McIver LJ, Blanco-Míguez A, Dubois L, Asnicar F, Maharjan S, Mailyan A, Manghi P, Scholz M, Thomas AM, et al. Integrating taxonomic, functional, and strain-level profiling of diverse microbial communities with bioBakery 3. Elife 2021; 10:e65088.

47. Buchfink B, Xie C, Huson DH. Fast and sensitive protein alignment using DIAMOND. Nat Methods 2015; 12:59–60.

48. The UniProt Consortium. UniProt: the universal protein knowledgebase. Nucleic Acids Res 2017; 45:D158–69.

49. Lex A, Gehlenborg N, Strobelt H, Vuillemot R, Pfister H. UpSet: visualization of intersecting sets. IEEE Trans Vis Comput Graph 2014; 20:1983–92.

50. Dixon P. VEGAN, a package of R functions for community ecology. Journal of Vegetation Science 2003; 14:927–30.

51. Gu Z, Eils R, Schlesner M. Complex heatmaps reveal patterns and correlations in multidimensional genomic data. Bioinformatics 2016; 32:2847–9.

52. Swertz MA, Dijkstra M, Adamusiak T, van der Velde JK, Kanterakis A, Roos ET, Lops J, Thorisson GA, Arends D, Byelas G, et al. The MOLGENIS toolkit: rapid prototyping of biosoftware at the push of a button. BMC Bioinformatics 2010; 11 Suppl 12:S12.

53. Signorell A, Aho K, Alfons A, Anderegg N, Aragon T, Arachchige C, Arppe A, Baddeley A, Barton K, Bolker B, et al. DescTools: tools for descriptive statistics [Internet]. 2024; Available from: https://cran.r-project.org/web/packages/DescTools/index.html

54. Maechler M, Rousseeuw P, Struyf A, Hubert M, Hornik K, Studer M, Roudier P, Gonzalez J, Kozlowski K, Schubert E, et al. cluster: “Finding groups in data”: cluster analysis extended Rousseeuw et al. [Internet]. 2023; Available from: https://cran.r-project.org/web/packages/cluster/index.html

55. Hennig C. fpc: Flexible Procedures for Clustering [Internet]. 2024; Available from: https://cran.r-project.org/web/packages/fpc/index.html

56. Kuznetsova A, Brockhoff PB, Christensen RHB, Jensen SP. lmerTest: tests in linear mixed effects models [Internet]. 2020; Available from: https://cran.r-project.org/web/packages/lmerTest/index.html

57. Schippa S, Iebba V, Barbato M, Di Nardo G, Totino V, Checchi MP, Longhi C, Maiella G, Cucchiara S, Conte MP. A distinctive “microbial signature” in celiac pediatric patients. BMC Microbiol 2010; 10:175.

58. Pisani A, Rausch P, Bang C, Ellul S, Tabone T, Marantidis Cordina C, Zahra G, Franke A, Ellul P. Dysbiosis in the gut microbiota in patients with inflammatory bowel disease during remission. Microbiology Spectrum 2022; 10:e00616–22.

59. Vila AV, Hu S, Andreu-Sánchez S, Collij V, Jansen BH, Augustijn HE, Bolte LA, Ruigrok RAAA, Abu-Ali G, Giallourakis C, et al. Original research: Faecal metabolome and its determinants in inflammatory bowel disease. Gut 2023; 72:1472.

60. Bourgonje AR, Roo-Brand G, Lisotto P, Sadaghian Sadabad M, Reitsema RD, de Goffau MC, Faber KN, Dijkstra G, Harmsen HJM. Patients with inflammatory bowel disease show IgG immune responses towards specific intestinal bacterial genera. Front Immunol 2022; 13:842911.

61. Pittayanon R, Lau JT, Yuan Y, Leontiadis GI, Tse F, Surette M, Moayyedi P. Gut microbiota in patients with irritable bowel syndrome—a systematic review. Gastroenterology 2019; 157:97–108.

62. Di Cagno R, De Angelis M, De Pasquale I, Ndagijimana M, Vernocchi P, Ricciuti P, Gagliardi F, Laghi L, Crecchio C, Guerzoni ME, et al. Duodenal and faecal microbiota of celiac children: molecular, phenotype and metabolome characterization. BMC Microbiol 2011; 11:219.

63. El Mouzan M, Al-Hussaini A, Serena G, Assiri A, Al Sarkhy A, Al Mofarreh M, Alasmi M, Fasano A. Microbiota profile of new-onset celiac disease in children in Saudi Arabia. Gut Pathog 2022; 14:37.

64. Sánchez E, Laparra JM, Sanz Y. Discerning the role of *Bacteroides fragilis* in celiac disease pathogenesis. Appl Environ Microbiol 2012; 78:6507–15.

65. Sánchez E, Donat E, Ribes-Koninckx C, Calabuig M, Sanz Y. Intestinal *Bacteroides* species associated with coeliac disease. Journal of Clinical Pathology 2010; 63:1105–11.

66. García-López M, Meier-Kolthoff JP, Tindall BJ, Gronow S, Woyke T, Kyrpides NC, Hahnke RL, Göker M. Analysis of 1,000 type-strain genomes improves taxonomic classification of *Bacteroidetes*. Front Microbiol 2019; 10:2083.

67. De Palma G, Nadal I, Collado MC, Sanz Y. Effects of a gluten-free diet on gut microbiota and immune function in healthy adult human subjects. Br J Nutr 2009; 102:1154–60.

68. Valitutti F, Cucchiara S, Fasano A. Celiac disease and the microbiome. Nutrients 2019; 11:2403.

69. Lionetti E, Dominijanni V, Iasevoli M, Cimadamore E, Acquaviva I, Gatti S, Monachesi C, Catassi G, Pino A, Faragalli A, et al. Effects of the supplementation with a multispecies probiotic on clinical and laboratory recovery of children with newly diagnosed celiac disease: A randomized, placebo-controlled trial. Digestive and Liver Disease 2023; 55:1328–37.

70. Dieterich W, Schuppan D, Schink M, Schwappacher R, Wirtz S, Agaimy A, Neurath MF, Zopf Y. Influence of low FODMAP and gluten-free diets on disease activity and intestinal microbiota in patients with non-celiac gluten sensitivity. Clin Nutr 2019; 38:697–707.

71. Bonder MJ, Tigchelaar EF, Cai X, Trynka G, Cenit MC, Hrdlickova B, Zhong H, Vatanen T, Gevers D, Wijmenga C, et al. The influence of a short-term gluten-free diet on the human gut microbiome. Genome Med 2016; 8:45.

72. Rosero JA, Killer J, Sechovcová H, Mrázek J, Benada O, Fliegerová K, Havlík J, Kopečný J. Reclassification of *Eubacterium rectale* (Hauduroy et al. 1937) Prévot 1938 in a new genus *Agathobacter* gen. nov. as *Agathobacter rectalis* comb. nov., and description of *Agathobacter ruminis* sp. nov., isolated from the rumen contents of sheep and cows. Int J Syst Evol Microbiol 2016; 66:768–73.

73. Sheridan PO, Duncan SH, Walker AW, Scott KP, Louis P, Flint HJ. Objections to the proposed reclassification of *Eubacterium rectale* as *Agathobacter rectalis*. Int J Syst Evol Microbiol 2016; 66:2106.

74. Rosero JA, Killer J, Sechovcová H, Mrázek J, Benada O, Fliegerová K, Havlík J, Kopečný J. Reply to the Letter to the Editor by Paul O. Sheridan, Sylvia H. Duncan, Alan W. Walker, Karen P. Scott, Petra Louis and Harry J. Flint, referring to our paper “Reclassification of *Eubacterium rectale* (Prévot et al. 1967) in a new genus *Agathobacter gen. nov.*, as *Agathobacter rectalis comb. nov.*, within the family Lachnospiraceae, and description of *Agathobacter ruminis sp. nov.*, from the rumen”, Int J Syst Evol Microbiol, DOI 10.1099/ijsem.0.000788. Int J Syst Evol Microbiol 2016; 66:2107.

75. Zuo G, Hao B. Whole-genome-based phylogeny supports the objections against the reclassification of *Eubacterium rectale* to *Agathobacter rectalis*. Int J Syst Evol Microbiol 2016; 66:2451.

76. Gophna U, Konikoff T, Nielsen HB. *Oscillospira* and related bacteria – From metagenomic species to metabolic features. Environmental Microbiology 2017; 19:835–41.

77. Karcher N, Pasolli E, Asnicar F, Huang KD, Tett A, Manara S, Armanini F, Bain D, Duncan SH, Louis P, et al. Analysis of 1321 *Eubacterium rectale* genomes from metagenomes uncovers complex phylogeographic population structure and subspecies functional adaptations. Genome Biol 2020; 21:138.

78. Ruiz L, Bottacini F, Boinett CJ, Cain AK, O’Connell-Motherway M, Lawley TD, van Sinderen D. The essential genomic landscape of the commensal *Bifidobacterium breve* UCC2003. Sci Rep 2017; 7:5648.

79. Iyer LM, Zhang D, Rogozin IB, Aravind L. Evolution of the deaminase fold and multiple origins of eukaryotic editing and mutagenic nucleic acid deaminases from bacterial toxin systems. Nucleic Acids Res 2011; 39:9473–97.

80. Amiri P, Hosseini SA, Ghaffari S, Tutunchi H, Ghaffari S, Mosharkesh E, Asghari S, Roshanravan N. Role of butyrate, a gut microbiota derived metabolite, in cardiovascular diseases: a comprehensive narrative review. Front Pharmacol 2021; 12:837509.

81. Liu H, Wang J, He T, Becker S, Zhang G, Li D, Ma X. Butyrate: a double-edged sword for health? Adv Nutr 2018; 9:21–9.

82. Singh V, Lee G, Son H, Koh H, Kim ES, Unno T, Shin J-H. Butyrate producers, “The Sentinel of Gut”: Their intestinal significance with and beyond butyrate, and prospective use as microbial therapeutics. Front Microbiol 2022; 13:1103836.

83. Baldi S, Menicatti M, Nannini G, Niccolai E, Russo E, Ricci F, Pallecchi M, Romano F, Pedone M, Poli G, et al. Free fatty acids signature in human intestinal disorders: significant association between butyric acid and celiac disease. Nutrients 2021; 13:742.

84. Tjellström B, Högberg L, Stenhammar L, Fälth-Magnusson K, Magnusson K-E, Norin E, Sundqvist T, Midtvedt T. Faecal short-chain fatty acid pattern in childhood coeliac disease is normalised after more than one year’s gluten-free diet. Microb Ecol Health Dis 2013; 24:10.3402/mehd.v24i0.20905.

85. Lionetti E, Antonucci N, Marinelli M, Bartolomei B, Franceschini E, Gatti S, Catassi GN, Verma AK, Monachesi C, Catassi C. Nutritional status, dietary intake, and adherence to the mediterranean diet of children with celiac disease on a gluten-free diet: a case-control prospective study. Nutrients 2020; 12:143.

86. Rinninella E, Cintoni M, Raoul P, Triarico S, Dionisi T, Gasbarrini GB, Gasbarrini A, Mele MC. The healthy gluten-free diet: practical tips to prevent metabolic disorders and nutritional deficiencies in celiac patients. Gastroenterology Insights 2021; 12:166–82.

87. Vici G, Belli L, Biondi M, Polzonetti V. Gluten free diet and nutrient deficiencies: A review. Clin Nutr 2016; 35:1236–41.

88. De Palma G, Nadal I, Medina M, Donat E, Ribes-Koninckx C, Calabuig M, Sanz Y. Intestinal dysbiosis and reduced immunoglobulin-coated bacteria associated with coeliac disease in children. BMC Microbiol 2010; 10:63.

89. Sánchez E, De Palma G, Capilla A, Nova E, Pozo T, Castillejo G, Varea V, Marcos A, Garrote JA, Polanco I, et al. Influence of environmental and genetic factors linked to celiac disease risk on infant gut colonization by *Bacteroides* species. Appl Environ Microbiol 2011; 77:5316–23.

90. De Palma G, Capilla A, Nova E, Castillejo G, Varea V, Pozo T, Garrote JA, Polanco I, López A, Ribes-Koninckx C, et al. Influence of milk-feeding type and genetic risk of developing coeliac disease on intestinal microbiota of infants: the PROFICEL study. PLoS One 2012; 7:e30791.

91. Esteban-Torres M, Ruiz L, Rossini V, Nally K, van Sinderen D. Intracellular glycogen accumulation by human gut commensals as a niche adaptation trait. Gut Microbes 15:2235067.

92. Brüssow H. Problems with the concept of gut microbiota dysbiosis. Microb Biotechnol 2019; 13:423–34.

93. Tojo R, Suárez A, Clemente MG, de los Reyes-Gavilán CG, Margolles A, Gueimonde M, Ruas-Madiedo P. Intestinal microbiota in health and disease: role of bifidobacteria in gut homeostasis. World J Gastroenterol 2014; 20:15163–76.

94. Ferretti G, Bacchetti T, Masciangelo S, Saturni L. Celiac disease, inflammation and oxidative damage: a nutrigenetic approach. Nutrients 2012; 4:243–57.

95. Ramírez-Sánchez AD, Zühlke S, Aguirre-Gamboa R, Vochteloo M, Franke L, Lundin KEA, Withoff S, Jonkers IH. Gene expression and eQTL analysis reflect the heterogeneity in the inflammatory status of the duodenal epithelial lining in coeliac disease. 2024; :2024.02.29.582756.

96. Galhardo RS, Hastings PJ, Rosenberg SM. Mutation as a stress response and the regulation of evolvability. Critical Reviews in Biochemistry and Molecular Biology 2007; 42:399.

97. Radman M. SOS repair hypothesis: phenomenology of an inducible DNA repair which is accompanied by mutagenesis. Basic Life Sci 1975; 5A:355–67.

98. Prudhomme M, Attaiech L, Sanchez G, Martin B, Claverys J-P. Antibiotic stress induces genetic transformability in the human pathogen *Streptococcus pneumoniae*. Science 2006; 313:89–92.

99. Slager J, Kjos M, Attaiech L, Veening J-W. Antibiotic-induced replication stress triggers bacterial competence by increasing gene dosage near the origin. Cell 2014; 157:395–406.

100. Richie TG, Wiechman H, Ingold C, Heeren L, Kamke A, Pogranichniy S, Monk K, Summers T, Ran Q, Sarkar S, et al. Eubacterium rectale detoxification mechanism increases resilience of the gut environment [Internet]. 2024; :2024.05.09.593360. Available from: https://www.biorxiv.org/content/10.1101/2024.05.09.593360v1

101. Bourgonje AR, Andreu-Sánchez S, Vogl T, Hu S, Vich Vila A, Gacesa R, Leviatan S, Kurilshikov A, Klompus S, Kalka IN, et al. Phage-display immunoprecipitation sequencing of the antibody epitope repertoire in inflammatory bowel disease reveals distinct antibody signatures. Immunity 2023; 56:1393–1409.e6.

102. Lodes MJ, Cong Y, Elson CO, Mohamath R, Landers CJ, Targan SR, Fort M, Hershberg RM. Bacterial flagellin is a dominant antigen in Crohn disease. J Clin Invest 2004; 113:1296–306.

103. Hagopian WA, Lernmark A, Rewers MJ, Simell OG, She J-X, Ziegler AG, Krischer JP, Akolkar B. TEDDY--The Environmental Determinants of Diabetes in the Young: an observational clinical trial. Ann N Y Acad Sci 2006; 1079:320–6.

104. Eurén A, Lynch K, Lindfors K, Parikh H, Koletzko S, Liu E, Akolkar B, Hagopian W, Krischer J, Rewers M, et al. Risk of celiac disease autoimmunity is modified by interactions between CD247 and environmental exposures. Sci Rep 2024; 14:25463.

105. Lindfors K, Lin J, Lee H-S, Hyöty H, Nykter M, Kurppa K, Liu E, Koletzko S, Rewers M, Hagopian W, et al. Metagenomics of the faecal virome indicate a cumulative effect of enterovirus and gluten amount on the risk of coeliac disease autoimmunity in genetically at risk children: the TEDDY study. Gut 2020; 69:1416–22.

