## Supplementary Figures S1-S4 for "High-resolution analysis of the treated coeliac disease microbiome reveals strain-level variation"

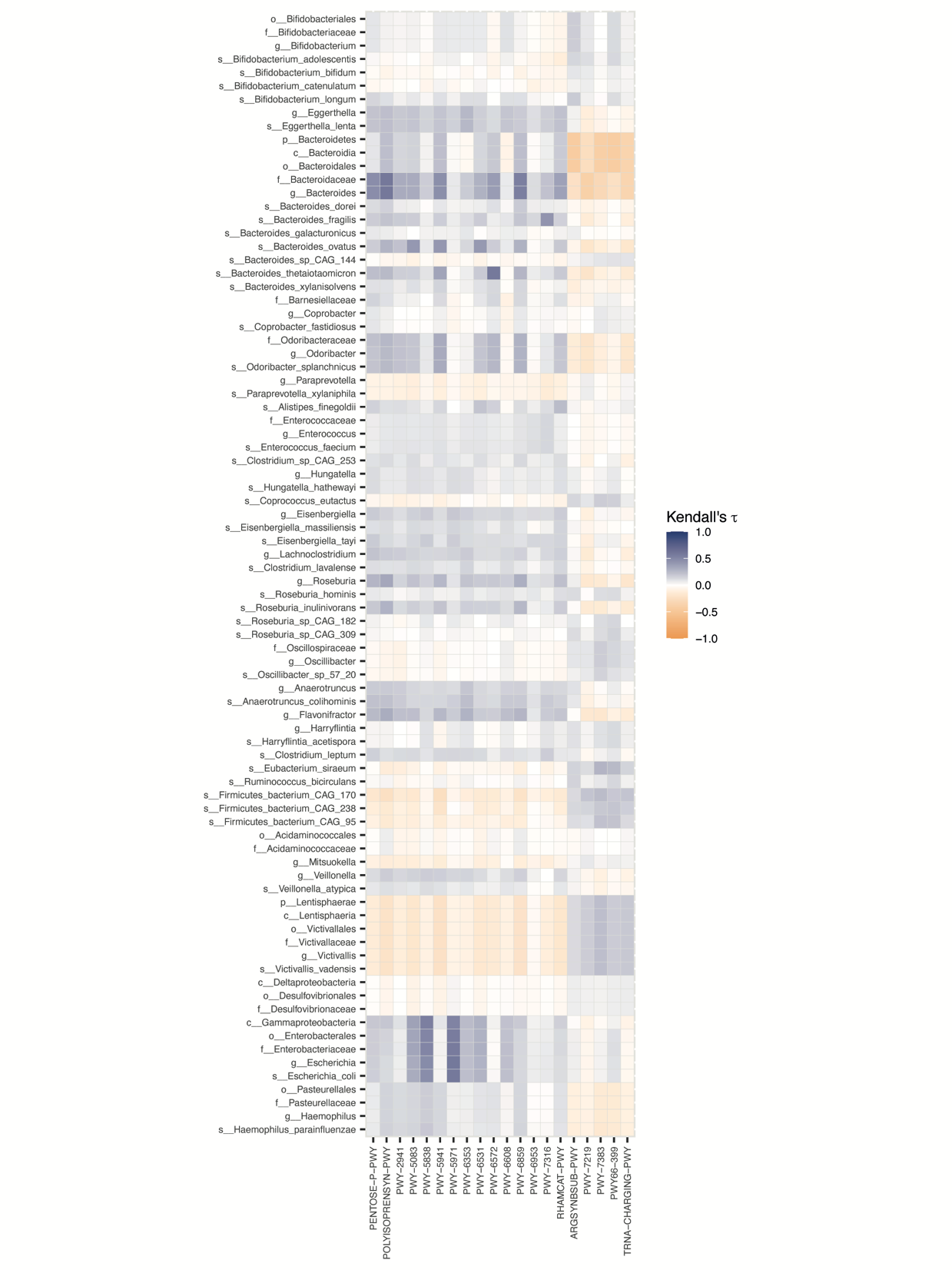

Supp. Figure S1: Correlation matrix between differentially abundant metabolic pathways (columns) and bacterial taxa (rows). The genus *Bacteroides* and higher-level taxa are included for visualization purposes.

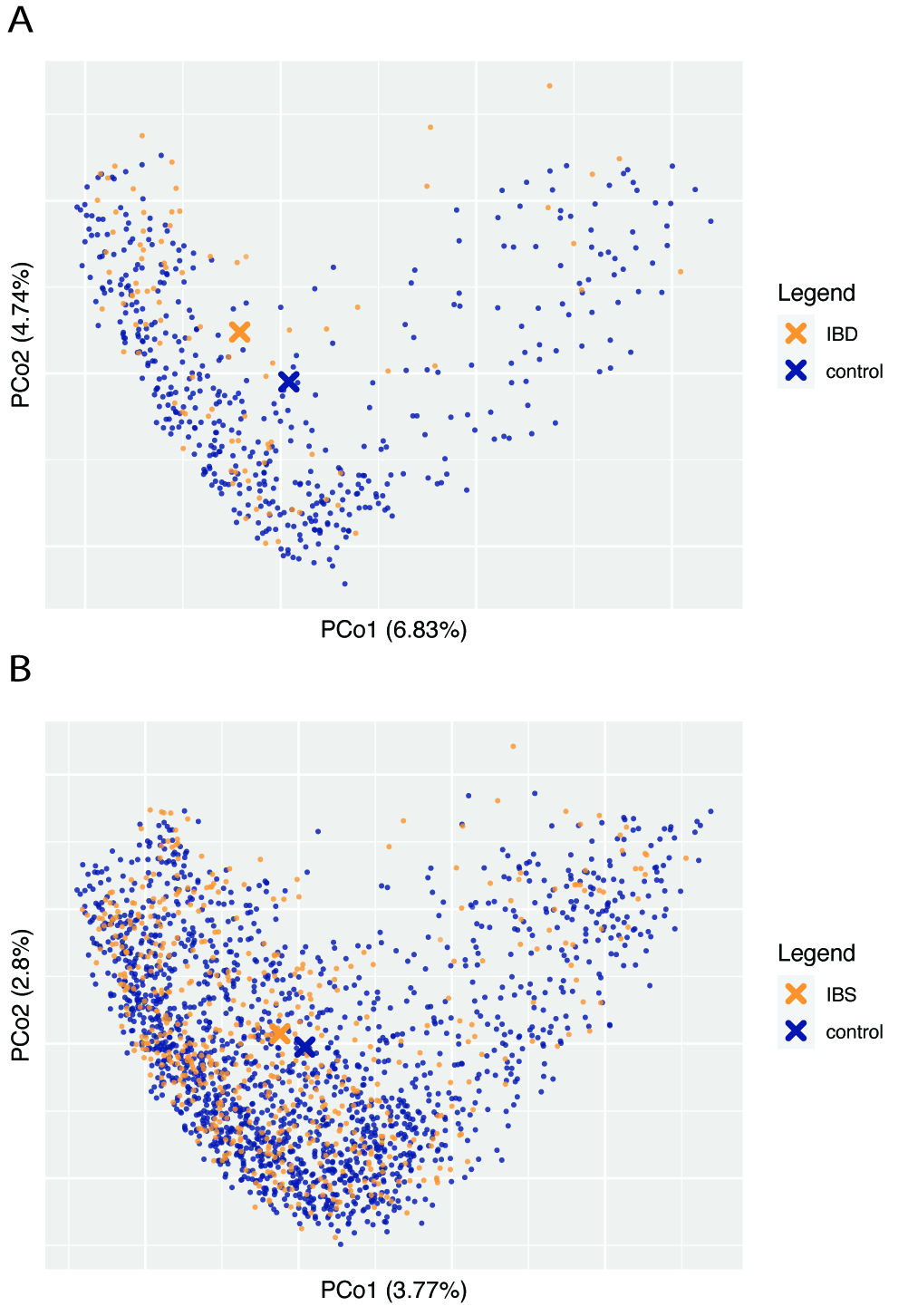

Supp. Figure S2: Beta diversity in IBD and IBS vs. controls, using Bray-Curtis dissimilarity as a distance metric. A. Principal coordinate plot of IBD samples and matched controls. Centroid locations are indicated with a cross. B. Principal coordinate plot of IBS samples and matched controls. Centroid locations are indicated with a cross.

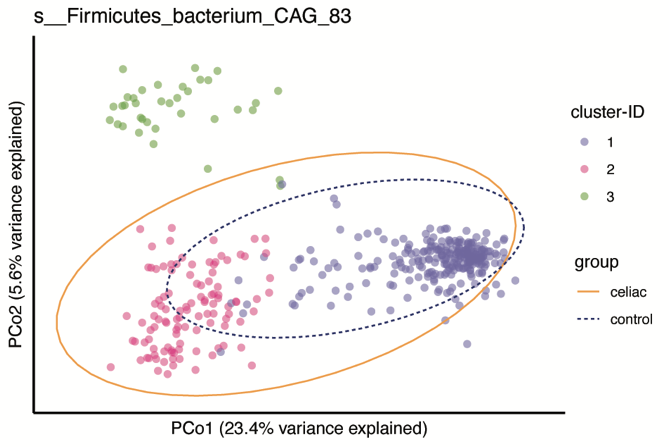

Supp. Figure S3: Non-metric multidimensional scaling plots of pairwise genetic distances between *Firmicutes bacterium CAG:83* strains. Points are colored according to their assigned cluster (PAM clustering). Ellipses include 95% of the samples in the indicated group.

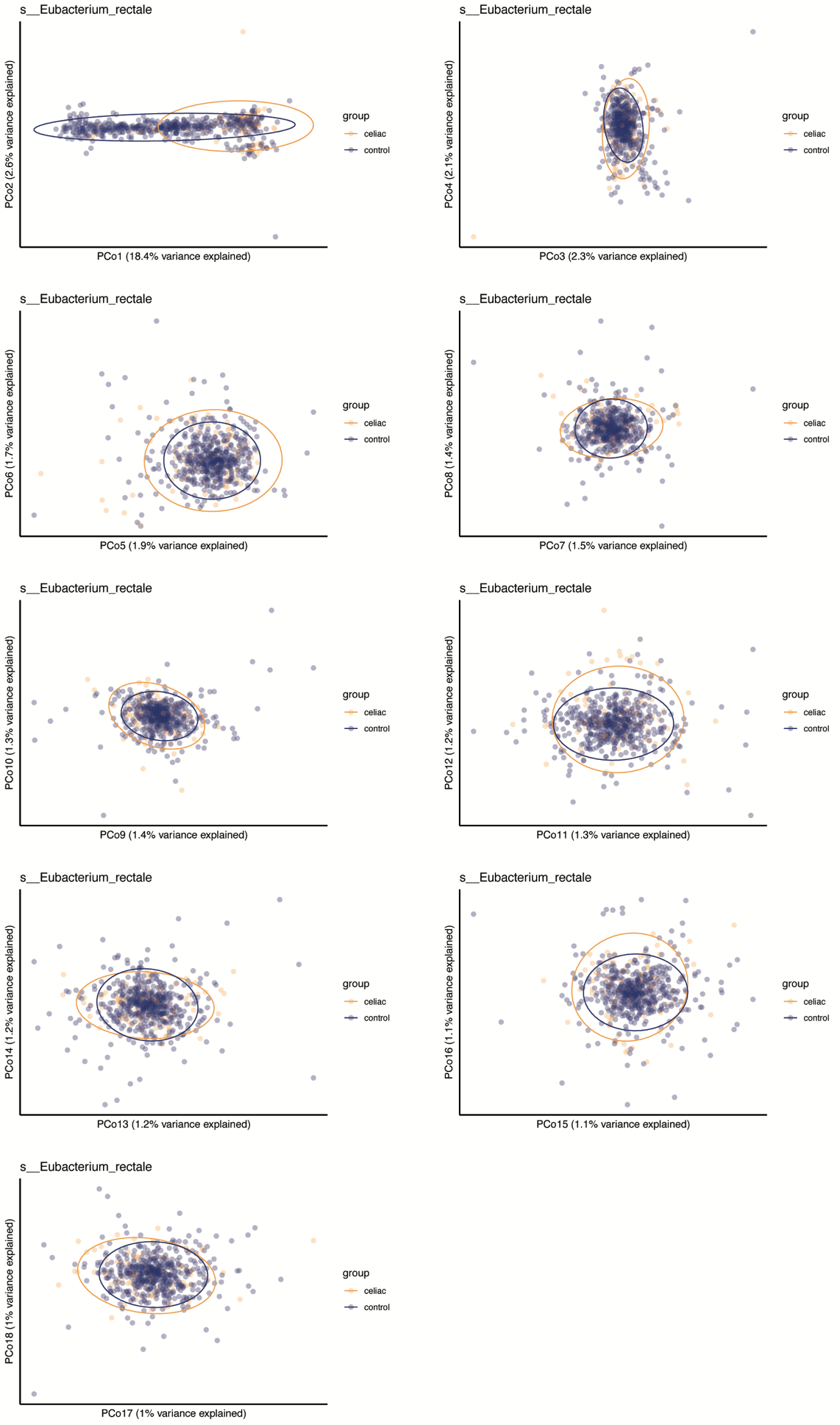

Supp. Figure S4: Non-metric multidimensional scaling plots of pairwise genetic distances between *E. rectale* strains, colored by disease status. All principal coordinates are shown that explain more than 1% of variation among samples. Ellipses include 95% of the samples in the indicated group.
